# *EARLY FLOWERING 3* controls temperature responsiveness of the circadian clock

**DOI:** 10.1101/2020.11.11.378307

**Authors:** Zihao Zhu, Marcel Quint, Muhammad Usman Anwer

**Author notes:** **Author Contributions** M.U.A. and M.Q. conceived the project. Z. Z. performed the experiments. All authors contributed to write the article. **Funding information** The funding for this work was provided by grants from the European Social Fund and the Federal State of Saxony-Anhalt (International Graduate School AGRIPOLY - Determinants of Plant Performance, grant ZS/2016/08/80644) and by the Deutsche Forschungsgemeinschaft (Qu 141/6-1) to MQ.

## Abstract

Predictable changes in light and temperature during a diurnal cycle are major entrainment cues that enable the circadian clock to generate internal biological rhythms that are synchronized with the external environment. With the average global temperature predicted to keep increasing, the intricate light-temperature coordination that is necessary for clock functionality is expected to be seriously affected. Hence, understanding how temperature signals are perceived by the circadian clock has become an important issue, especially in light of climate change scenarios. In *Arabidopsis*, the clock component *EARLY FLOWERING 3* (*ELF3*) not only serves as an essential light *Zeitnehmer*, but also functions as a thermosensor participating in thermomorphogenesis. However, the role of *ELF3* in temperature entrainment of the circadian clock is not fully understood. Here, we report that *ELF3* is essential for delivering temperature input to the clock. We demonstrate that in the absence of *ELF3*, the oscillator was unable to properly respond to temperature changes, resulting in an impaired gating of thermoresponses. Consequently, clock-controlled physiological processes such as rhythmic growth and cotyledon movement were disturbed. Together, our results reveal that *ELF3* is an essential *Zeitnehmer* for temperature sensing of the oscillator, and thereby for coordinating the rhythmic control of thermoresponsive physiological outputs.

## Introduction

While latitudinal daylength remains stable, global warming results in increasing ambient temperatures. As a consequence, the intrinsic system that relies on the integrated daylength-temperature signals to control important physiological and developmental outputs could be seriously affected (Schaarschmidt *et al.*, 2020). This can lead to abnormal growth and developmental patterns that potentially result in serious yield losses in important crops (Quint *et al.*, 2016; Lippmann *et al.*, 2019). Since this trend of global temperature increase is predicted to continue, it is important to understand how key regulatory networks perceive light and temperature signals to control fundamental developmental and physiological processes.

The circadian clock is one such endogenous key network that utilizes external cues (also known as *Zeitgeber*), primarily light and temperature, as timing input to precisely synchronize internal cellular mechanisms with the external environment. The timing information from the *Zeitgeber* is received by oscillator components known as *Zeitnehmer* that help to reset and synchronize the clock with the external environment. This *Zeitgeber-Zeitnehmer* communication is called entrainment that subsequently sets the pace of the oscillator (Oakenfull & Davis, 2017). Once entrained, the oscillators generate a ~24h rhythmicity that can be sustained for long periods; even in the absence of environmental cues (i.e., free-running conditions, such as constant light and temperature). After synchronizing with the external environment, oscillators regulate the rhythmic accumulation of several transcripts, proteins, and metabolites. The circadian clock thereby confers fitness advantages by allowing organisms to anticipate and adapt to the changing environment (Covington *et al.*, 2008; Yamashino *et al.*, 2008; Anwer *et al.*, 2014; Ronald & Davis, 2017).

The central part of the clock, the oscillator, is composed of transcriptional-translational feedback loops (TTFLs) (Nohales & Kay, 2016). In *Arabidopsis thaliana* (Arabidopsis) three such loops, a morning loop, an evening loop and a central loop, constitute the oscillator. The central loop is a dual negative feedback loop comprised of two partially redundant MYB-like transcription factors CIRCADIAN CLOCK ASSOCIATED 1 (CCA1) and LATE ELONGATED HYPOCOTYL (LHY), and a member of the PSEUDO-RESPONSE REGULATOR (PRR) family TIMING OF CAB EXPRESSION 1 (TOC1/PRR1) (Alabadı́ *et al.*, 2001; Huang *et al.*, 2012), which negatively regulate each other’s expression. In the morning loop, CCA1/LHY repress *PRR7* and *PRR9,* which later repress *CCA1/LHY* (Nakamichi *et al.*, 2010; Adams *et al.*, 2015). The evening expression of TOC1 represses *GIGANTEA* (*GI*), which in turn activates *TOC1*, which together form the evening loop (Kim *et al.*, 2007; Huang *et al.*, 2012). Besides these three fundamental loops, a complex of three evening phased proteins (known as evening complex or EC), consisting of ELF4, ELF3 and LUX ARRYTHMO (LUX), has been established as an integral part of the oscillator. The EC directly represses the transcription of the morning loop member *PRR9* and the evening loop component *GI* (Nusinow *et al.*, 2011; Herrero *et al.*, 2012; Ezer *et al.*, 2017). Furthermore, CCA1 directly represses *ELF3* and thereby connects the EC with the central loop (Lu *et al.*, 2012; Kamioka *et al.*, 2016).

A constant *Zeitgeber-Zeitnehmer* communication in the entrainment process is critical to keep the oscillator in-phase with the external environment (Anwer *et al.*, 2020). The phytochrome B (phyB) photoreceptor functions as both light and temperature sensor and thereby is an important component of the entrainment mechanism. However, phyB does not act as *Zeitnehmer*, since it is neither required for clock entrainment, nor for oscillator function (Sanchez *et al.*, 2020). The interactions of phyB with ELF3 and GI present one possible *Zeitgeber-Zeitnehmer* junction through which light and temperature information may be delivered to the oscillator (Anwer *et al.*, 2020). Consistently, severe light and temperature signaling anomalies have been observed in *elf3* and *gi* mutants (Kolmos *et al.*, 2011; Anwer *et al.*, 2014; Panigrahi & Mishra, 2015; Anwer *et al.*, 2020). For instance, under free-running conditions oscillator defects such as arrhythmia in *elf3* and altered circadian periodicity in *gi* have been reported (Anwer *et al.*, 2014; Anwer *et al.*, 2020). Besides that, both mutants display several other pleiotropic phenotypes such as elongated hypocotyl and altered flowering time, suggesting that several important clock-regulated downstream pathways are also disrupted (McWatters *et al.*, 2000; Yamashino *et al.*, 2008; Kim *et al.*, 2012; Anwer *et al.*, 2014; Box *et al.*, 2015; Raschke *et al.*, 2015; Anwer *et al.*, 2020). Not surprisingly, they share common targets such as *PHYTOCHROME-INTERACTING FACTOR 4* (*PIF4*) and *FLOWERING LOCUS T* (*FT*) (Nusinow *et al.*, 2011; Anwer *et al.*, 2020) to regulate important physiological and developmental processes such as growth and flowering time, respectively.

We recently demonstrated that photoperiod-responsive growth and flowering time was lost in *elf3 gi* double mutants, and established that these two genes are essential for clock entrainment to light signals (Anwer *et al.*, 2020). However, the mechanism of clock entrainment in response to temperature cycles is still poorly understood (Avello *et al.*, 2019). Among the little that is known, *PRR7* and *PRR9* are two components with conceivable roles in the temperature input to the oscillator (Salomé & McClung, 2005). The *prr7 prr9* double mutant displayed conditional arrhythmia depending on the temperature regime used during the entrainment (Salomé & McClung, 2005; Salomé *et al.*, 2010). This suggests that these components are required for temperature input to the oscillator in a temperature-dependent manner.

The role of the EC (ELF3-ELF4-LUX) in the temperature input is also intriguing. All components of the EC physically bind to the recently established thermosensor phyB. Furthermore, the binding of the EC to its target gene promoters is temperature-dependent (Kolmos *et al.*, 2011; Herrero *et al.*, 2012; Kim *et al.*, 2013; Box *et al.*, 2015; Huang & Nusinow, 2016; Ezer *et al.*, 2017). Consistently, the EC has been proposed to be a night-time repressor of the temperature input to the clock (Mizuno *et al.*, 2014). This contradicts a previous finding that demonstrated *ELF3* to be an integral part of the oscillator and advocated against its function as a *Zeitnehmer* in the temperature-input pathway (Thines & Harmon, 2010). However, a recent finding that highlighted ELF3 as a temperature sensor (independently of the EC) re-emphasizes the need of a comprehensive study examining *ELF3* function in temperature entrainment (Jung *et al.*, 2020).

In this study, we systematically investigate the role of *ELF3* in temperature entrainment of the circadian clock. We demonstrate that *ELF3* is essential for temperature input to the oscillator. In the absence of *ELF3*, the circadian oscillator failed to respond and synchronize to external temperature cycles. Furthermore, our data demonstrate that *ELF3* is also fundamental for the clock gating ability, which is essential to generate rhythmic processes by precisely allowing temperature information to pass through only during an optimum time window within a diurnal cycle. Our data thus establish *ELF3* as an essential temperature *Zeitnehmer* in the circadian oscillator. In the scenario of global warming, this understanding may be helpful to improve crop performance under higher temperatures.

## Materials and methods

### Plant Materials and growth conditions

All *Arabidopsis thaliana* lines used were in the Ws-2 background. The *elf3-4*, *gi-158* and *elf3-4 gi-158* null mutants have been described previously (Zagotta *et al.*, 1996; Hicks *et al.*, 2001; Anwer *et al.*, 2020). Sterilized *Arabidopsis* seeds were cold stratified for 3 d in darkness, and were allowed to germinate on solid *Arabidopsis thaliana* solution (ATS) nutrient medium with 1% (weight : volume) sucrose (Lincoln *et al.*, 1990). Unless stated otherwise, seedlings were grown on vertically oriented plates in long day (LD, 16 h light : 8 h dark) or short day (SD, 8 h light : 16 h dark) with 90 μmol m^−2^s^−1^ photosynthetically active radiation (PAR) using white fluorescent lamps (T5 4000K). Seedlings were grown at constant 16°C or 22°C for 8 d, or at constant 20°C or 28°C for 8 d. For temperature shift assays, seedlings grown at 20°C for 4 d were shifted to 28°C or were kept at 20°C for additional 4 d. For assays in constant light (LL, 90 μmol m^−2^s^−1^), seedlings were grown at constant 16°C, 22°C or 28°C for 8 d. Seedlings were imaged, and hypocotyl length was measured using ImageJ (http://image.nih.gov/ij/).

### Measurements of growth rate and elevation angle

To allow unobstructed visualization of hypocotyl and cotyledons in air, seedlings were grown vertically on the agar ledge formed by removing part of the agar in the square plate as previously described (Anwer *et al.*, 2020). Imaging was started at Zeitgeber Time (ZT) 00 on day 3. Photographs were taken every 60 min for 96 h in constant light (LL, white fluorescent lamps: 30 μmol m^−2^s^−1^) under specified thermocycles (12 h 22°C : 12 h 16°C or 12 h 28°C : 12 h 22°C). For free-running conditions, seedlings were entrained by thermocycles for 2 d and then on day 3 at ZT00 were released into constant conditions (30 μmol m^−2^s^−1^ light and 22°C temperature). The imaging platform with infrared illumination was previously described (Anwer *et al.*, 2020). Image stacks were analyzed using ImageJ (http://image.nih.gov/ij/). The circadian parameters of cotyledon movement were determined using the MFourFit method integrated in the BioDare2 analysis platform (Zielinski *et al.*, 2014). The relative amplitude error (RAE) analysis was used to estimate the robustness of the circadian rhythm: RAE values range from 0 to 1, where 0 represents a robust rhythm, and 1 represents no rhythm.

### Analysis of transcript levels

Seedlings were entrained in constant light (LL, 90 μmol m^−2^s^−1^) or darkness (DD), under 12 h 22°C : 12 h 16°C thermocycles for 8 d. On day 9, starting from ZT00, the samples were harvested every 4 h. For the temperature-gating assay, seedlings were entrained under thermocycles (with LL) as described above for 8 d. On day 9, starting from ZT00, seedlings were either treated with a 4 h temperature pulse (28°C) at various ZTs, or were kept under same conditions (no treatment) before samples were harvested at the specified time. All experiments were performed using three biological replicates. Isolation of total RNA samples from whole seedlings, reverse transcription-mediated quantitative real-time polymerase chain reaction (RT-qPCR) and primer sequences have been described previously (Anwer *et al.*, 2020). The primers used for *PRR7* were forward: 5’-TGAAAGTTGGAAAAGGACCA-3’ and reverse: 5’-GTTCCACGTGCATTAGCTCT-3’.

## Results

### *ELF3* and *GI* are involved in temperature-photoperiod crosstalk

Temperature and light independently and collaboratively serve as two prominent entrainment cues of the circadian clock (Eckardt, 2005; Avello *et al.*, 2019; Gil & Park, 2019). The major red-light photoreceptor phyB that functions also as a thermosensor (Jung *et al.*, 2016; Legris *et al.*, 2016; Delker *et al.*, 2017) stabilizes ELF3 protein, suggesting a possible light-temperature signal transduction pathway (Reed *et al.*, 2000; Liu *et al.*, 2001; Nieto *et al.*, 2015). A previous study showed that *elf3* mutants were arrhythmic in continuous darkness (DD) after temperature entrainment (Thines & Harmon, 2010). However, since the phyB thermosensor is essentially nonfunctional in darkness, it remains unclear whether the role of *ELF3* in thermocycle entrainment is still sustained in light, with phyB activated. To investigate thermocycle entrainment in the presence of light, we decided to first estimate the extent of a possible temperature-photoperiod interconnection, since both *ELF3* and *GI* control the photoperiod sensing of the circadian clock (Anwer *et al.*, 2020). We used cellular elongation of the hypocotyl as a classic phenotypic readout, which is known to be highly responsive to both temperature and photoperiod variations (Niwa *et al.*, 2009). We measured the hypocotyl length of Ws-2, null mutants *elf3-4* and *gi-158,* and *elf3-4 gi-158* seedlings grown in long day (LD, 16 h light : 8 h dark, Fig.1a) or short day (SD, 8 h light : 16 h dark, Fig. 1b) conditions. To estimate temperature response under these photoperiods, the seedlings were grown at constant 16°C or 22°C for 8 d before hypocotyl measurements were taken. We found that the higher temperature resulted in the acceleration of growth in all four genotypes in both photoperiods (Fig. 1a,b). However, the extent of the response to higher temperature in LD or SD was different in these four genotypes. We found that Ws-2 and *gi-158* were more responsive in SD than in LD, whereas *elf3-4* and *elf3-4 gi-158* displayed the opposite result (Fig. 1c).

**Fig. 1.**
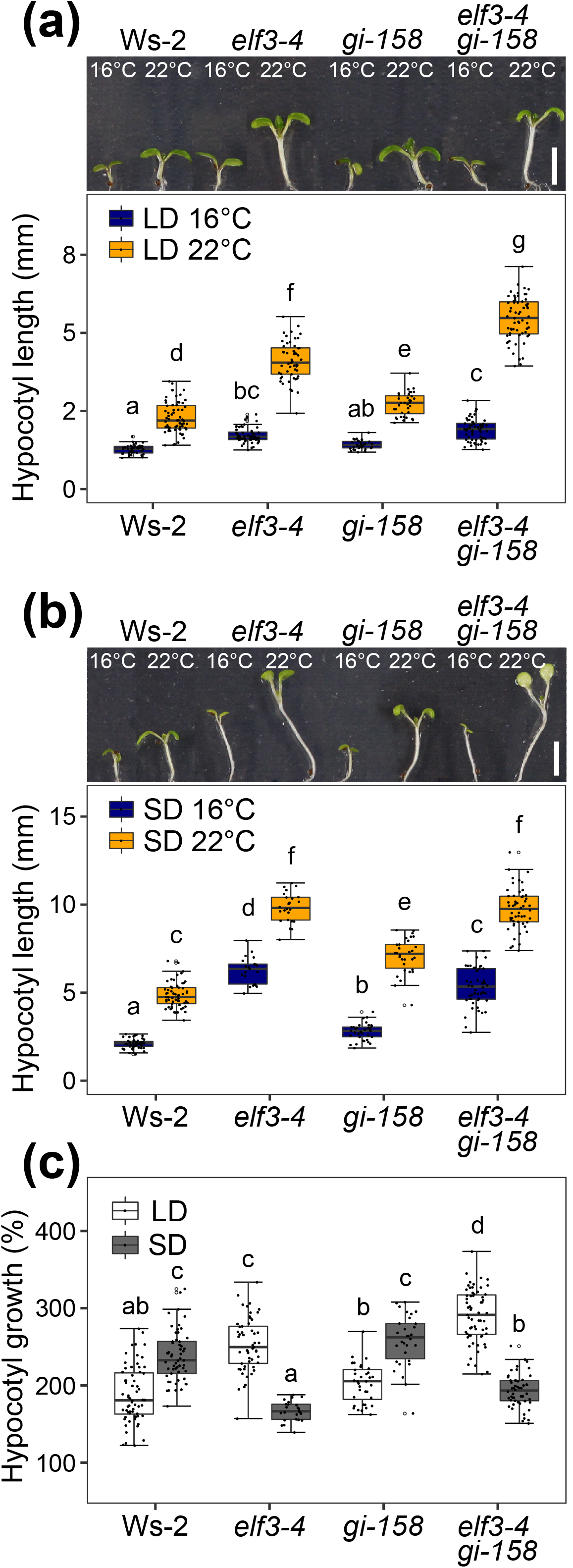
*ELF3* and *GI* are involved in temperature-photoperiod crosstalk. (a, b) Representative images and quantification of the hypocotyl length of 8-d-old Ws-2, *elf3-4*, *gi-158* and *elf3-4 gi-158* seedlings grown in LD (16 h light : 8 h dark, a) and SD (8 h light : 16 h dark, b). Seedlings were grown at constant 16°C or 22°C for 8 d. Scale bars = 4mm. (c) Hypocotyl length at 22°C relative to the median at 16°C as shown in (a) and (b). Box plots show medians and interquartile ranges. Dots represent biological replicates, and those greater than 1.5x interquartile range are outliers. Different letters above the boxes indicate significant differences (two-way ANOVA with Tukey’s HSD test, *P* < 0.05).

Similar results were also observed in seedlings grown under similar photoperiods, but at a higher temperature regime (constant 20°C or 28°C). Here, only Ws-2 was more responsive to temperature in SD (6 h light : 18 h dark, Fig. S1b) than in LD (18 h light : 6 h dark, Fig. S1a), whereas all three mutants displayed the opposite result (Fig. S1a-c). We detected a similar response in a temperature shift assay, where the 4-d-old seedlings grown at 20°C were shifted to 28°C or were kept at 20°C for additional 4 d before hypocotyl measurements were taken (Fig. S1d-f). In addition, *elf3-4 gi-158* double mutant displayed an additive effect on hypocotyl length in LD (Fig. S1a,d), but not in SD (Fig. 1b, Fig. S1b,e) or under constant 16°C (Fig. 1a,b). These data demonstrate that (i) *elf3* and *gi* mutants respond to ambient temperatures differently than the wild type Ws-2, and that (ii) temperature responsiveness of especially *elf3* but also *gi* mutants is strongly influenced by the photoperiod. Together, this suggests that *ELF3* and *GI* are important participants of a likely rather complicated temperature-photoperiod crosstalk.

Next, we sought to determine whether the thermoresponsive growth remains intact in the absence of photocycles and whether *ELF3* and *GI* play any significant role in determining the responsiveness to temperature under these non-cycling conditions. We examined the thermoresponsiveness of hypocotyl elongation in continuous light (LL), by measuring hypocotyl length of Ws-2, *elf3-4*, *gi-158*, and *elf3-4 gi-158* seedlings grown in LL at constant temperature of 16°C, 22°C or 28°C (Fig. S2). In contrast to the previous experiment, we found that in the absence of photocycles the temperature response of Ws-2 and all three mutant lines was largely similar (Fig. S2). As such, temperature response defects in *elf3* and *gi* mutants depend on the presence of photocycles, while their temperature response seems intact in the absence of photoperiods (LL). Taken together, our data indicate that both *ELF3* and *GI* play important roles in temperature-photoperiod crosstalk, however, they are not essential for temperature responsiveness under non-cycling conditions.

### Clock-controlled physiological processes require *ELF3* under thermocycles

The circadian clock controls rhythmic oscillation patterns of several physiological processes such as hypocotyl growth and leaf movement. Under diurnal conditions, circadian oscillators coordinate hypocotyl elongation with daily environmental changes such as photoperiod, resulting in maximum growth rate at dawn or early morning in SD and LD, respectively (Nozue *et al.*, 2007; Niwa *et al.*, 2009; Anwer *et al.*, 2020). This is largely processed by the growth-repressive function of *ELF3* and *GI* during the night and day times, respectively (Anwer *et al.*, 2020).

While the conclusions of the data shown so far (Fig. 1, Fig. S1, Fig. S2) apply to non-cycling temperature conditions, we now aimed to understand the role of *ELF3* and *GI* under cycling temperature conditions. To investigate whether and how *ELF3* and *GI* contribute to rhythmic hypocotyl elongation in seedlings under temperature cycles (hereafter thermocycles), we measured the growth rates of Ws-2, *elf3-4*, *gi-158* and *elf3-4 gi-158* every hour for 4 d under thermocycles (12 h 22°C : 12 h 16°C) in the absence of photocycles (LL) (Fig. 2a,b, Table S1). We used these conditions to circumvent potential temperature-photoperiod crosstalk as shown above (Fig. 1, Fig. S1).

**Fig. 2.**
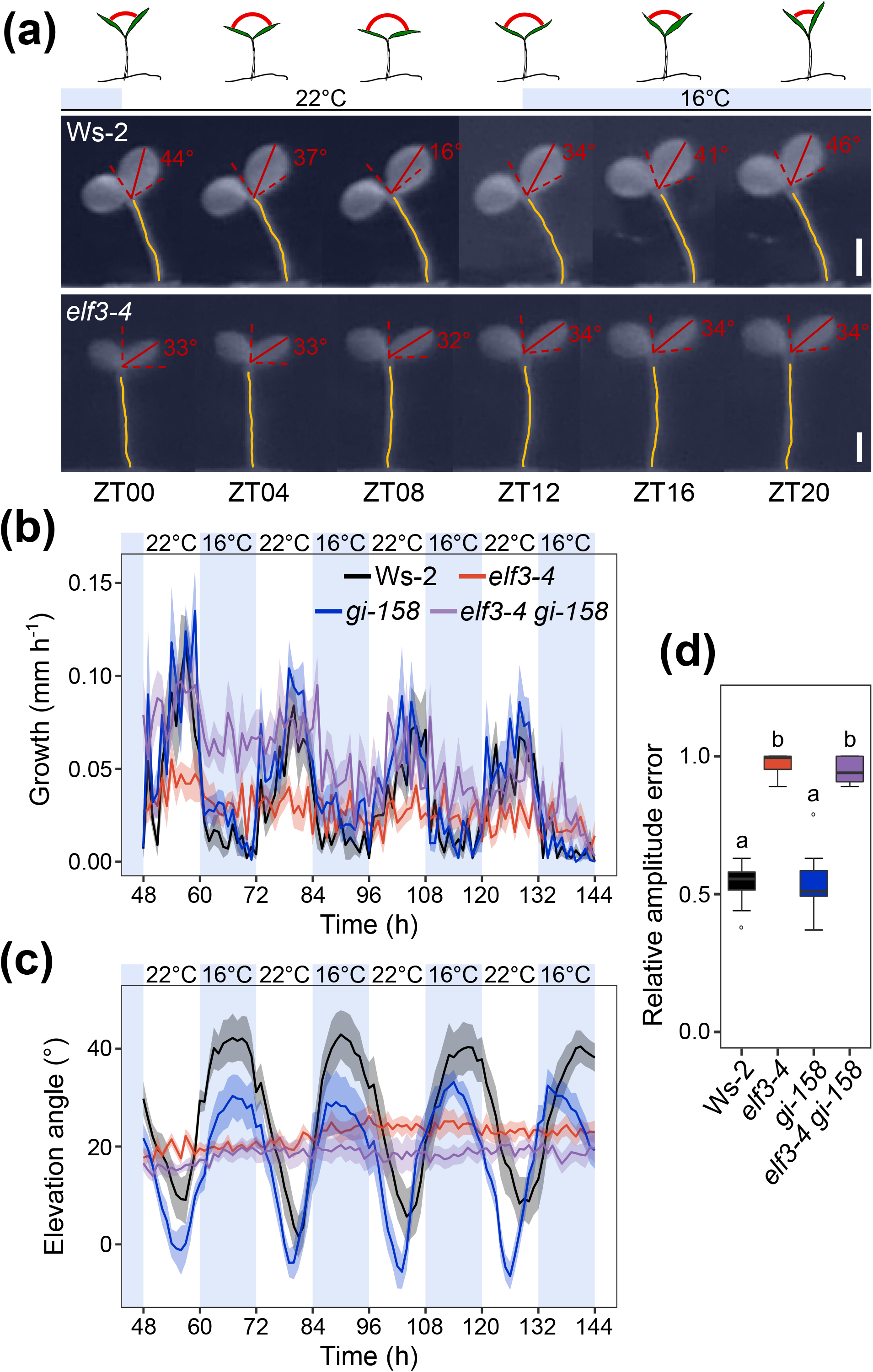
Rhythmic growth and cotyledon movement under thermocycles require *ELF3*. (a) Representative images of 5-d-old Ws-2 and *elf3-4* seedlings grown in LL under thermocycles. Non-shaded areas represent warm period (22°C), whereas blue-shaded areas represent cold period (16°C). Representative photographs taken every 4 h starting from ZT00 on day 6 are shown. Relative coordinates (dashed red lines) were generated to ensure that both cotyledons had the same position, no matter whether the position of the whole seedling changed or not during growth. The measured angles (physical quantities in red) between cotyledon position (from tip to base, red lines) and relative horizontal (dashed red lines in horizontal) are defined as elevation angles. The yellow lines indicate the measured length of the hypocotyls. Scale bars = 1 mm. Sketches above the images are shown for illustration purposes and represent the cotyledon movement of a hypothetical plant. The red arcs represent the hypothetical angles between two cotyledons. (b, c) Quantification of hypocotyl growth (b) and cotyledon elevation angle (c) of Ws-2, *elf3-4*, *gi-158* and *elf3-4 gi-158* seedlings, grown under thermocycles as in (a). Starting from ZT00 on day 3, photographs were taken every hour. Lines represent the mean and ribbons indicate standard error of mean (SEM) (n = 8). (d) Relative amplitude error of cotyledon movement data shown in (c). Box plots show medians and interquartile ranges. Outliers (greater than 1.5x interquartile range) are marked with open circles. Different letters above the boxes indicate significant differences (one-way ANOVA with Tukey’s HSD test, *P* < 0.05).

In Ws-2 and *gi-158*, we detected rhythmic growth patterns with maximum growth rates during mid to late stages (~ZT08) of the warm period (22°C) (Fig. 2b, Table S1). In contrast, no clear growth peaks were detected in *elf3-4* and *elf3-4 gi-158* (Fig. 2b, Table S1). In *elf3-4*, we detected a constant growth rate, which was much lower than Ws-2 during the warm period (22°C) and marginally higher during the cool period (16°C). In *elf3-4 gi-158*, the growth rates were similar to Ws-2 during the warm period (22°C), but were much higher during the cool period (16°C). Importantly, just like *elf3-4*, no clear growth peaks were detected (Fig. 2b, Table S1). Thus, these data indicated that rhythmic growth under thermocycles requires *ELF3*, while *GI* most likely only plays a minor role.

Like hypocotyl growth, cotyledon movement is another classic physiological output that is regulated by the circadian clock (Millar *et al.*, 1995). To further scrutinize the role of *ELF3* and *GI* in determining the functional capability of the clock under thermocycles, we measured the cotyledon elevation angle every hour of seedlings grown under thermocycles in LL (Fig. 2a,c). As expected for a functional clock, we detected rhythmic cotyledon movement in Ws-2 and *gi-158*, with open and closed cotyledons during the warm (22°C) and cool (16°C) periods, respectively (Fig. 2a,c). This is consistent with the previous report where similar patterns were observed in Col-0 and *gi-2* seedlings entrained by 12 h 22°C : 12 h 12°C thermocycles (Tseng *et al.*, 2004). However, in contrast to Ws-2 and *gi-158*, the cotyledon movement was undetectable in *elf3-4* and *elf3-4 gi-158* seedlings under the same conditions (Fig. 2a,c), mirroring the hypocotyl growth rate data (Fig. 2b, Table S1) and again suggesting a dysfunctional clock. The relative amplitude error (RAE) analysis confirmed robust rhythms of the cotyledon movement in Ws-2 and *gi-158* (RAE~0.5), whereas both *elf3-4* and *elf3-4 gi-158* were arrhythmic (RAE~1.0) (Fig. 2d).

To exclude the possibility that the rhythmic cotyledon movement observed in Ws-2 and *gi-158* was driven by the temperature variations rather than the circadian oscillator, the 2-d-thermocycle-entrained seedlings were transferred into free-running conditions (LL and constant 22°C) and cotyledon movement was measured (Fig. S3a). Consistent with the results under thermocycles, we detected robust rhythms in Ws-2 and *gi-158*, whereas both *elf3-4* and *elf3-4 gi-158* were arrhythmic in free-running conditions (Fig. S3a,b). Previous studies have reported a thermocycle-dependent arrhythmia in *prr7 prr9* double mutant, with *prr7 prr9* displaying robust rhythms under high-regime thermocycles (28°C : 22°C) and arrhythmia under low-regime thermocycles (22°C : 12°C) (Salomé & McClung, 2005; Salomé *et al.*, 2010). To investigate whether the observed arrhythmia in *elf3-4* and *elf3-4 gi-158* is also depending on the thermocycle temperature regime, we monitored the cotyledon movement of the Ws-2, *elf3-4*, *gi-158* and *elf3-4 gi-158* seedlings under high-regime thermocycles (12 h 28°C : 12 h 22°C, Fig. S3c,d) and also under free-running conditions (LL and constant 22°C, Fig. S3e,f) after high-regime thermocycle entrainment. Consistent with the low-regime thermocycle results (Fig. 2c,d), robust rhythms were not detected in *elf3-4* and *elf3-4 gi-158* (Fig. S3c-f), which were evident from high RAE values. Collectively, these data demonstrate that in contrast to clock-controlled rhythmic processes under photocycles (Anwer *et al.*, 2020), only *ELF3*, but not *GI*, is essential for clock-controlled rhythmic processes under thermocycles.

### The oscillator’s responsiveness to temperature changes requires *ELF3*

As rhythmic hypocotyl elongation and cotyledon movement are regulated by the circadian oscillator, we hypothesized that in the absence of *ELF3*, the central oscillator itself is dysfunctional in responding to temperature changes. To test this, we monitored the expression of the key central oscillator genes *CCA1*, *LHY*, *PRR9*, *PRR7* and *TOC1* under thermocycles in LL (Fig. 3a-e, Table S2). As expected for a functional oscillator, Ws-2 and *gi-158* showed rhythmic expression of these genes albeit differences in the expression levels were occasionally detected (Fig, 3a-e, Table S2). In Ws-2 and *gi-158*, *CCA1* and *LHY* displayed expression peaks at ZT00/24, *PRR9* at ZT04, *PRR7* at ZT08, and *TOC1* at ZT16 (Ws-2) or ZT12 (*gi-158*) (Fig. 3a-e, Table S2). In contrast, no rhythmic expression was detected in *elf3-4* and *elf3-4 gi-158* (Fig. 3a-e, Table S2). In these two mutants, almost no expression of *CCA1* and *LHY* can be detected, whereas *PRR9*, *PRR7* and *TOC1* maintained high levels of expression without oscillations (Fig. 3a-e, Table S2). Similar patterns of rhythmic gene expression were also detected when plants were grown under the same thermocycles in darkness (Fig. S4a-e, Table S3). The exceptions were that, in DD, *gi-158* displayed an advanced expression peak of *PRR7* at ZT04 (Fig. S4d, Table S3), and slight peaks of *CCA1*, *LHY* and *PRR9* expression at ZT00/24 were also detected in *elf3-4* (Fig. S4a,b,c, Table S3). No clear expression pattern of *TOC1* was detected in all four genotypes under these conditions (Fig. S4e, Table S3). Together, these results indicate that *ELF3* is required to correctly set the phase of the key central oscillator genes in response to diurnal temperature changes.

**Fig. 3.**
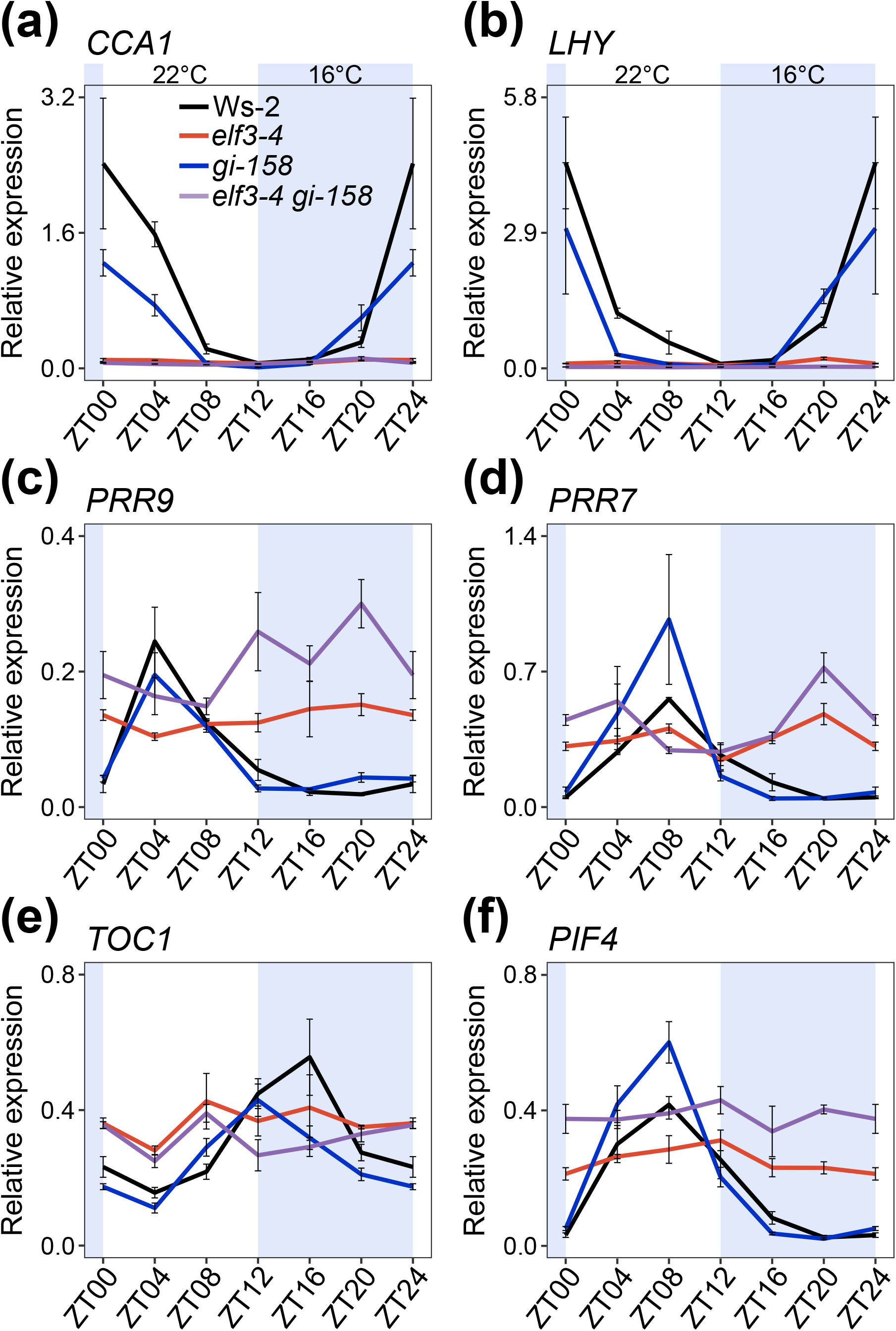
*ELF3* is required for oscillator’s responsiveness to temperature changes. (a-f) Transcript dynamics of key clock oscillator genes *CCA1* (a), *LHY* (b), *PRR9* (c), *PRR7* (d), *TOC1* (e) and major growth promoter *PIF4* (f). Ws-2, *elf3-4*, *gi-158* and *elf3-4 gi-158* seedlings were grown in LL under thermocycles for 8 d. On day 9, starting from ZT00, samples were harvested every 4 h. Non-shaded areas represent warm period (22°C), whereas blue-shaded areas represent cold period (16°C). Expression levels were normalized to *PROTEIN 19 PHOSPHATASE 2a subunit A3* (*PP2A*). Error bars indicate SEM (n = 3) of three biological replicates. The experiment was repeated twice with similar results.

As the circadian clock regulates thermoresponsive growth by regulating the major growth promoter *PIF4* (Nusinow *et al.*, 2011; Box *et al.*, 2015; Raschke *et al.*, 2015), we next monitored the expression of *PIF4* as a proxy to gauge the oscillator’s ability to regulate its target genes under thermocycles in LL (Fig. 3f, Table S2) and DD (Fig. S4f, Table S3). We found that, in Ws-2 and *gi-158*, the expression of *PIF4* specifically peaked during the warm period (22°C) at ZT08 in LL (Fig. 3f, Table S2), consistent with their rhythmic hypocotyl elongation (Fig. 2b, Table S1). In DD, Ws-2 displayed the *PIF4* expression peak at the same time (ZT08) as in LL, whereas *PIF4* expression was advanced and peaked at ZT04 in *gi-158* (Fig. S4f, Table S2). Importantly, in contrast to Ws-2 and *gi-158*, no clear peak of *PIF4* expression was detected in *elf3-4* and *elf3-4 gi-158*. Both displayed pronounced high *PIF4* expression, especially during the cool period (16°C) in both LL (Fig. 3f, Table S2) and DD (Fig. S4f, Table S3). In *elf3-4* and *elf3-4 gi-158*, the *PIF4* expression remained the same at almost all time-points (Fig. 3f, Fig. S4f, Table S2, Table S3). Taken together, our data demonstrate that the oscillator’s ability to properly respond to temperature input depends on functional *ELF3*.

### *ELF3* is essential for precise gating of temperature signals

One hallmark property of the circadian clock is a mechanism called ‘gating’, in which the oscillator regulates its own sensitivity to environmental inputs such as light and temperature in a time-of-day dependent manner. This ensures that the downstream processes are not influenced by these environmental inputs in an untimely manner. For instance, a sudden change in light and temperature caused by a cloud covering the sun would have no substantial affect on the clock-controlled rhythmic processes. This gating process thereby plays fundamental role to maintain correct rhythms of the clock-controlled outputs. To test the clock’s gating ability in response to temperature, we monitored the expression of the key clock-regulated temperature-responsive genes *PRR7*, *PRR9* and *PIF4* in Ws-2, *elf3-4*, *gi-158* and *elf3-4 gi-158* (Fig. 4a,b, Fig. S5).

**Fig. 4.**
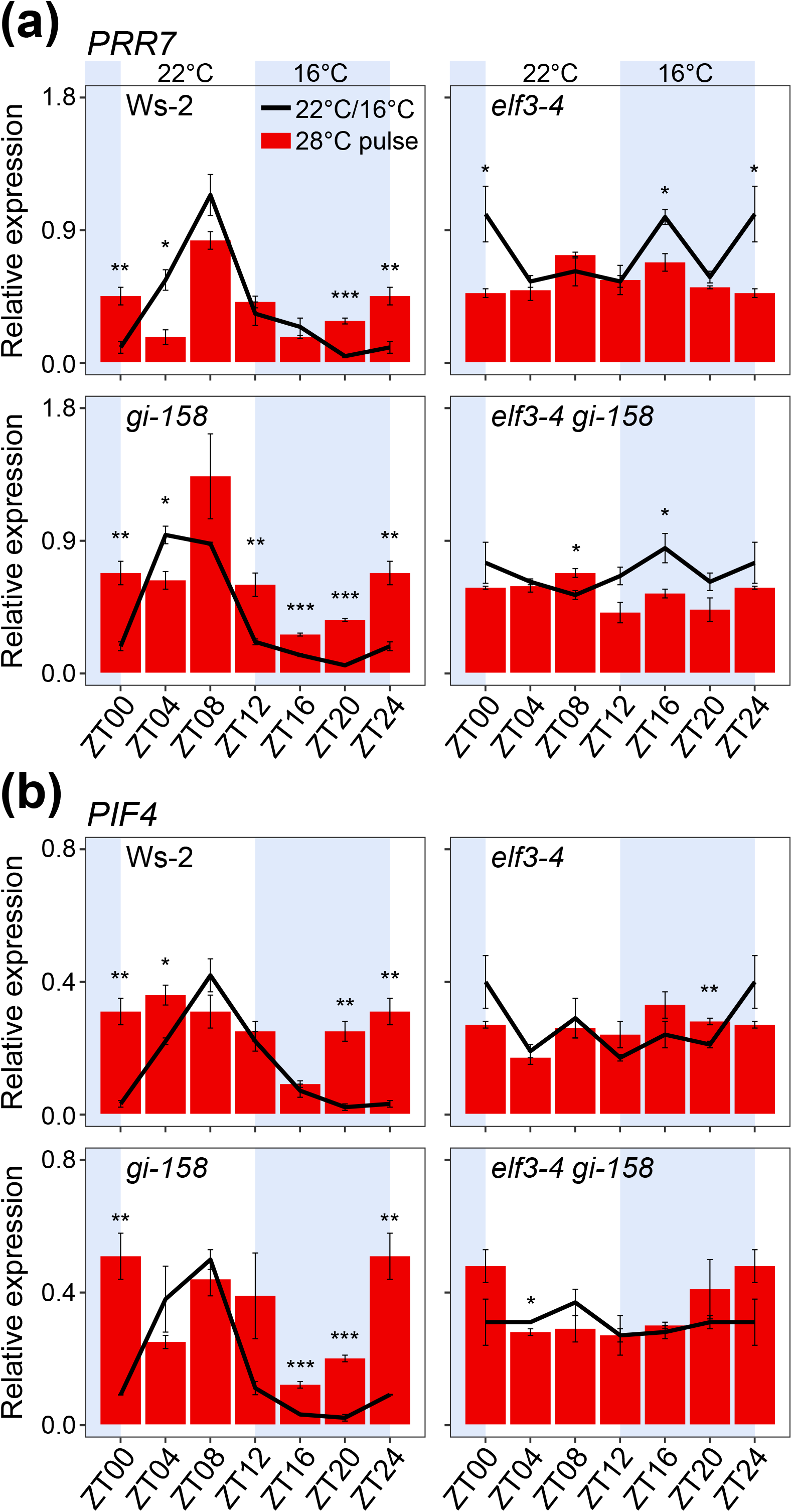
*ELF3* is essential for precise gating of the temperature signals under thermocycles. (a, b) Effect of the temperature pulse at the specified ZTs on the expression of *PRR7* (a) and *PIF4* (b). Ws-2, *elf3-4*, *gi-158* and *elf3-4 gi-158* seedlings were grown in LL under thermocycles for 8 d. On day 9, the seedlings were either treated with a 4 h temperature pulse (28°C pulse) at indicated ZTs, or were kept under same conditions (no treatment, 22°C/16°C) before samples were harvested. At indicated ZTs, red bars represent gene expression levels after treatment with a temperature pulse, whereas black lines represent gene expression levels at the same time without treatment. Non-shaded areas represent warm period (22°C), whereas blue-shaded areas represent cold period (16°C). Expression levels were normalized to *PP2A*. Error bars indicate SEM (n = 3) of three biological replicates. Asterisks above lines or bars indicate significant differences (*, *P* < 0.05; **, *P* < 0.01; ***, *P* < 0.001; Student’s *t*-test).

Thermocycle-entrained seedlings were either treated with a 4 h temperature pulse (28°C pulse) at various ZTs, or were kept under same conditions (no treatment) before samples were harvested at the specified time-points. We observed that in Ws-2, the temperature responsiveness of these genes was mainly restricted from late night to early morning (between ZT16-ZT04), as an induction of *PRR7*, *PRR9* and *PIF4* expression was detected primarily at these time-points (Fig. 4a,b, Fig. S5). In *gi-158*, the gates were opened slightly early, as an early induction of *PRR7* (ZT12-ZT24), and *PIF4* (ZT16-ZT24) was observed (Fig. 4a,b). Interestingly, the gating ability of the oscillator was abolished in *elf3-4* and *elf3-4 gi-158*. Except for some random time-points where the high-temperature response was opposite to the WT, mostly no response to temperature pulse was detected. Hence, the expression levels of *PRR7*, *PRR9* and *PIF4* remained unchanged at the vast majority of time-points (Fig. 4a,b, Fig. S5).

Taken together, these data demonstrate that *ELF3* is not only essential to generate robust rhythms under thermocycles but is also pivotal to maintain proper phase by blocking non-resetting temperature cues.

## Discussion

Increase in night-time ambient températures due to global warming could severely affect key regulatory mechanisms such as the circadian clock that relies on predictable changes in daily temperature cycles to coordinate essential biological events with the external environment (Schaarschmidt *et al.*, 2020). How the circadian clock utilizes diurnal temperature information to synchronize internal cellular mechanisms to the external environment remains largely unresolved. Here, we demonstrate that the circadian clock component *ELF3* is essential to establish communication between the circadian clock and ambient temperature. In the absence of *ELF3*, the circadian clock fails to respond to regular temperature cycles, which result in arrhythmia of key physiological processes (Fig. 2, Fig. 3, Fig. S3, Fig. S4, Table S1, Table S2). Thus, our data establish *ELF3* as a *Zeitnehmer* essential to relay temperature information to the circadian oscillator.

The involvement of *ELF3* and *GI* in light signaling has been reported since their identification (Zagotta *et al.*, 1996; Fowler *et al.*, 1999; Huq *et al.*, 2000; McWatters *et al.*, 2000; Kim *et al.*, 2007; Kolmos *et al.*, 2011). However, only recently we could conclusively show that both are necessary for clock entrainment to light cycles (Anwer *et al.*, 2020). It is important to note that the oscillator response to diurnal light signals remained intact in the absence of either *elf3-4* or *gi-158*. The oscillator only became non-responsive to photocycles when both components were absent (Anwer *et al.*, 2020). This is in contrast to the findings we report here for temperature entrainment, demonstrating that the oscillator’s ability to perceive temperature input during thermocycles is dependent largely on *ELF3* with *GI* playing only a minor role, if at all. Interestingly, besides these differences, we also observed a similar additive/synergistic relationship between *ELF3* and *GI* for thermocycles as reported for photocycles (Anwer et al., 2020). The hyperelongated hypocotyl under LD and LL (Fig. 1a, Fig. S1a,d, Fig. S2, Fig. S6), increased growth rate under thermocycles (Fig. 2b, Table S1), and overall higher expression of several genes (Fig. 3, Fig. S4, Table S2, Table S3) in *elf3-4 gi-158*, all consolidate their additive/synergistic function. However, clock entrainment to thermocycles is mainly dependent on *ELF3*.

In the literature, the role of *ELF3* as a temperature *Zeitnehmer* remained controversial. Thines and Harmon (2010) initially proposed that *ELF3* is an essential component of the oscillator but that it does not function as a *Zeitnehmer*. Their conclusions were based on the experiments performed on etiolated seedlings entrained to thermocycles in the darkness. Since under these conditions, phyB - a recently discovered temperature sensor that physically interacts with ELF3 - was absent (Jung *et al.*, 2016), the non-responsiveness of the oscillator to thermocycles could be partly attributed to the absence of phyB, leaving a major flaw in their study (which the authors could not have known back then). A later study then attempted to address these deficiencies by utilizing different photoperiod-temperature combinations for entrainment and highlighted the role of the EC in temperature input to the clock (Mizuno *et al.*, 2014). However, with a complicated cross-talk that exists between temperature and photoperiod (Fig. 1, Fig. S1) (Park *et al.*, 2020), it was hard to gauge the exact role of the *ELF3* in temperature entrainment.

Using thermocycles in constant light enabled us to eliminate these complications while maintaining the phyB-thermosensor activity. Under these conditions we here demonstrate that the circadian clock fails to entrain to thermocycles in the absence of *ELF3* (Fig. 2). Furthermore, proper responsiveness of the oscillator components to regular temperature changes (Fig. 3) as well as to sudden temperature pulses were also absent in the *elf3-4* mutant (Fig. 4, Fig. S5). Consequently, clock-controlled physiological processes such as cotyledon movement and diurnal hypocotyl growth were arrhythmic under thermocycles in *elf3-4* (Fig. 2, Fig. S3). Moreover, in confirmation of Thines and Harmon (2010), the *elf3-4* mutant failed to generate robust rhythms of key clock genes under thermocycles in darkness (Fig. S4). These data clearly indicate that *ELF3* is an essential *Zeitnehmer* that is pivotal for clock entrainment to temperature cycles. In conjunction with the recent finding that a prion-like domain in ELF3 functions as thermosensor, the necessity of phyB to mediate the temperature input to the oscillator can theoretically be excluded. However, the role of phyB in clock-independent thermomorphogenesis could not be eliminated since all four genotypes tested displayed robust temperature-responsive hypocotyl growth in constant light under non-cycling conditions (Fig. S2).

Contrary to the essential role of *ELF3* in temperature input to the clock, it is not fundamentally required to relay light signals to the oscillator. The *elf3-4* mutant, albeit with altered amplitude, was capable of generating robust rhythms under photocycles, indicating a partially functional oscillator (Anwer *et al.*, 2020). Such rhythms were entirely absent in the same mutant under thermocycles, highlighting the importance of *ELF3* in temperature responsiveness of the oscillator (Figs 2–4, Figs S3–S5). However, the loss of *GI* - another component of light signaling - along with *ELF3* absence resulted in similar clock dysfunction under photocycles (Anwer et al., 2020) as we demonstrate here for *elf3-4* under thermocycles. It therefore seems that the regulation of photocycle entrainment is more complex than the regulation of thermocycle entrainment. The existence of at least one additional component in the oscillator to maintain functionality under photocycles could be explained in an evolutionary context. First, under natural conditions, diurnal changes in light are the primary cue from which plants derive timing information. In principle, diurnal changes in temperature are just a byproduct of the presence or absence of the light. Second, being predominant photoautotrophs, plants require light to synthesize their food. Thus, on the one hand the presence of a robust clock provides a fitness advantage, on the other hand, however, a dysfunctional oscillator could be an existential threat. Therefore, not surprisingly, evolution has favored to develop a redundant mechanism to ensure a functional oscillator under light cycles.

Due to global warming, the intricate light-temperature relationship that is key to circadian clock functionality is being threatened by both relatively sudden as well as gradual increases in temperature (Lippmann *et al.*, 2019). Especially elevated temperature during the night (in the absence of light), which has already shown to affect crop yields, could seriously affect the clock’s ability to synchronize the internal biology with the external environment (Schaarschmidt *et al.*, 2020). Our data establish *ELF3* as an essential *Zeitnehmer* and provide mechanistic explanation of how temperature cues are perceived and processed by the circadian clock. Since *ELF3* is a known breeding target in key crops (Faure *et al.*, 2012; Bendix *et al.*, 2015), these findings provide insightful information to plant breeders to develop future crops which are more resilient to temperature changes.

---

**Fig. S1.**
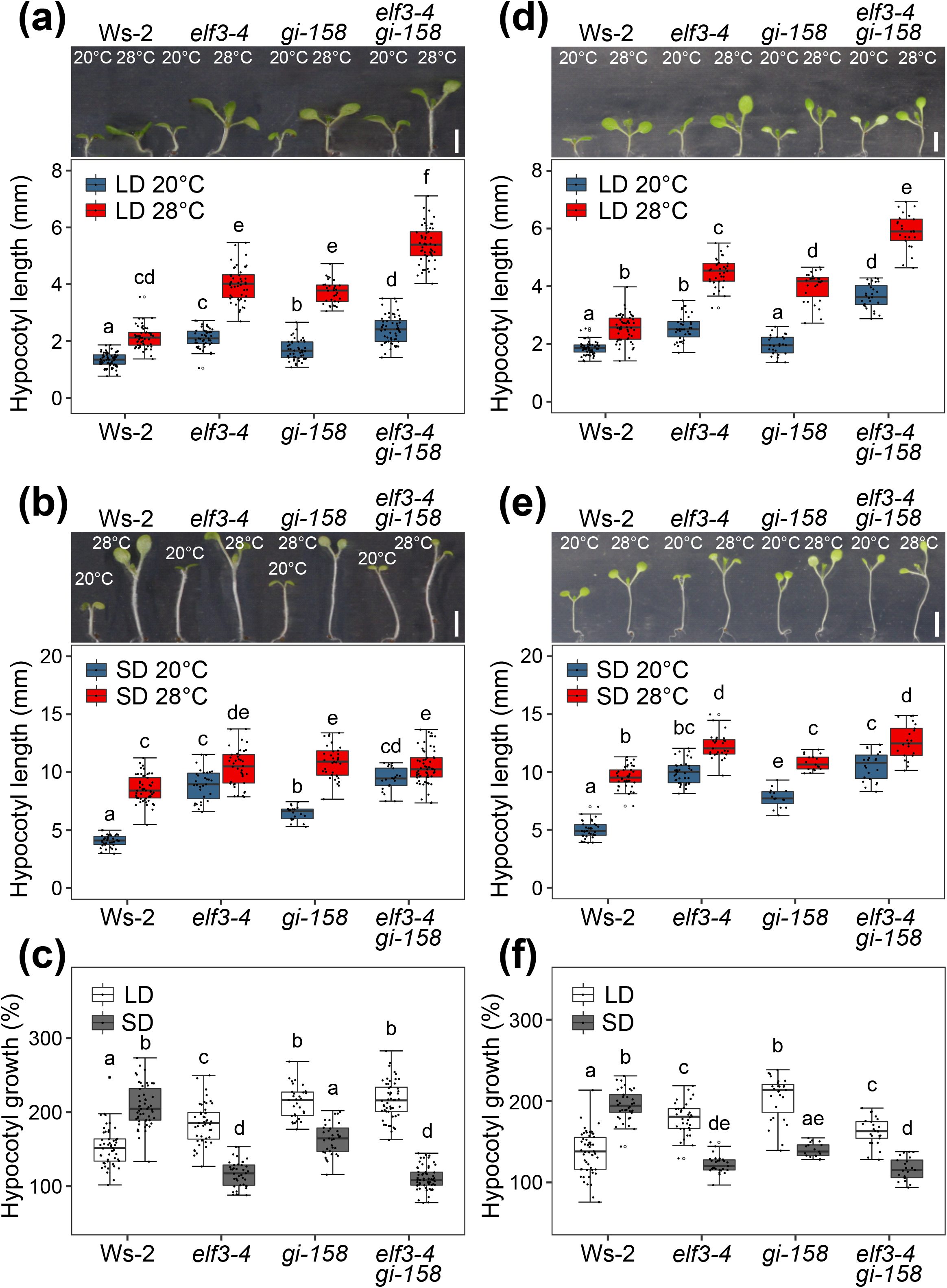
*ELF3* and *GI* are involved in temperature-photoperiod crosstalk at a higher temperature regime. (a, b, d, e) Representative images and quantification of the hypocotyl length of 8-d-old Ws-2, *elf3-4*, *gi-158* and *elf3-4 gi-158* seedlings grown in LD (18 h light : 6 h dark, a, d) and SD (6 h light : 18 h dark, b, e). (a, b) Seedlings were grown at constant 20°C or 28°C for 8 d. (d, e) Seedlings grown at 20°C for 4 d were shifted to 28°C or were kept at 20°C for additional 4 d. Scale bars = 4mm. (c, f) Hypocotyl length at 28°C relative to the median at 20°C as shown in (a) and (b), or in (d) and (e). Box plots show medians and interquartile ranges. Dots represent biological replicates, and those greater than 1.5x interquartile range are outliers. Different letters above the boxes indicate significant differences (two-way ANOVA with Tukey’s HSD test, *P* < 0.05).

**Fig. S2.**
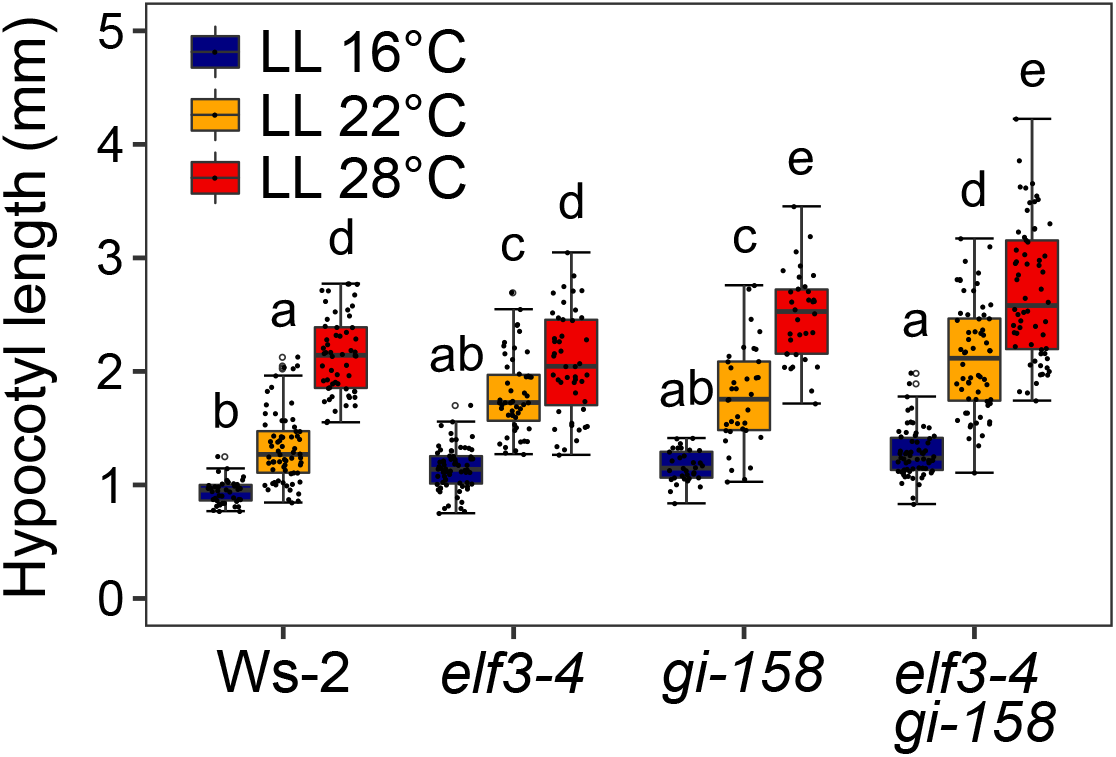
Thermoresponsive growth is intact in constant light. Quantification of the hypocotyl length of 8-d-old Ws-2, *elf3-4*, *gi-158* and *elf3-4 gi-158* seedlings grown in LL. Seedlings were grown at constant 16°C, 22°C or 28°C for 8 d. Box plots show medians and interquartile ranges. Dots represent biological replicates, and those greater than 1.5x interquartile range are outliers. Different letters above the boxes indicate significant differences (two-way ANOVA with Tukey’s HSD test, *P* < 0.05).

**Fig. S3.**
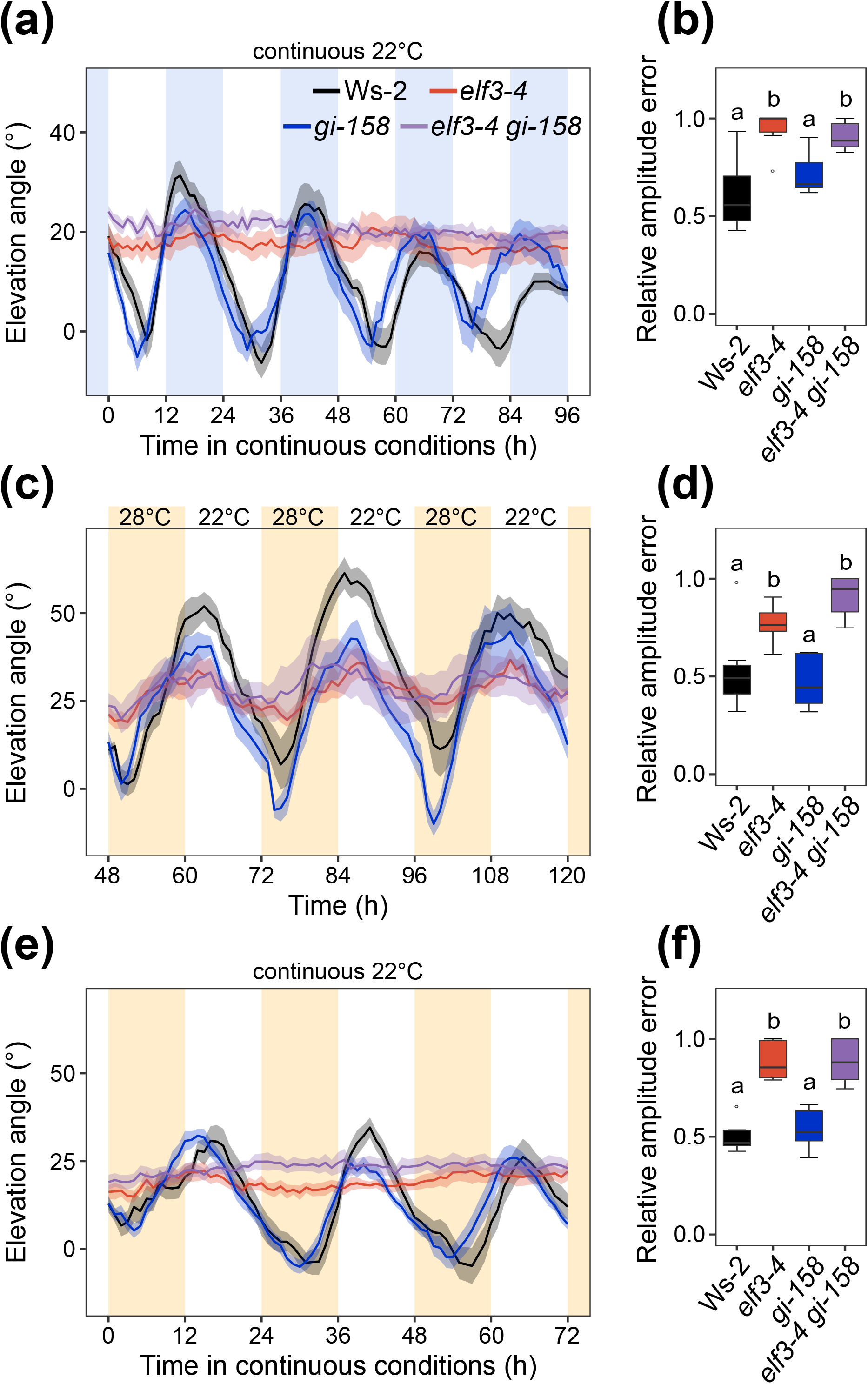
Rhythmic cotyledon movement under thermocycles requires *ELF3*. (a, c, e) Quantification of cotyledon elevation angle of Ws-2, *elf3-4*, *gi-158* and *elf3-4 gi-158* seedlings. (a) Seedlings were grown in LL under thermocycles for 2 d. On day 3, starting from ZT00, seedlings were released into constant conditions (LL and 22°C) and photographs were taken every hour. Non-shaded areas represent warm period (22°C), whereas blue-shaded areas represent cold period (16°C). (c) Seedlings were grown in LL under high-regime thermocycles for 2 d. On day 3, starting from ZT00, photographs were taken every hour. Orange-shaded areas represent warm period (28°C), whereas non-shaded areas represent cold period (22°C). (e) Seedlings were grown in LL under thermocycles as in (c). On day 3, starting from ZT00, seedlings were released into constant conditions (LL and 22°C) and photographs were taken every hour. Lines represent the mean and ribbons indicate SEM (n = 8). (b, d, f) Relative amplitude error of cotyledon movement data shown in (a), (c) and (e). Box plots show medians and interquartile ranges. Outliers (greater than 1.5x interquartile range) are marked with open circles. Different letters above the boxes indicate significant differences (one-way ANOVA with Tukey’s HSD test, *P* < 0.05).

**Fig. S4.**
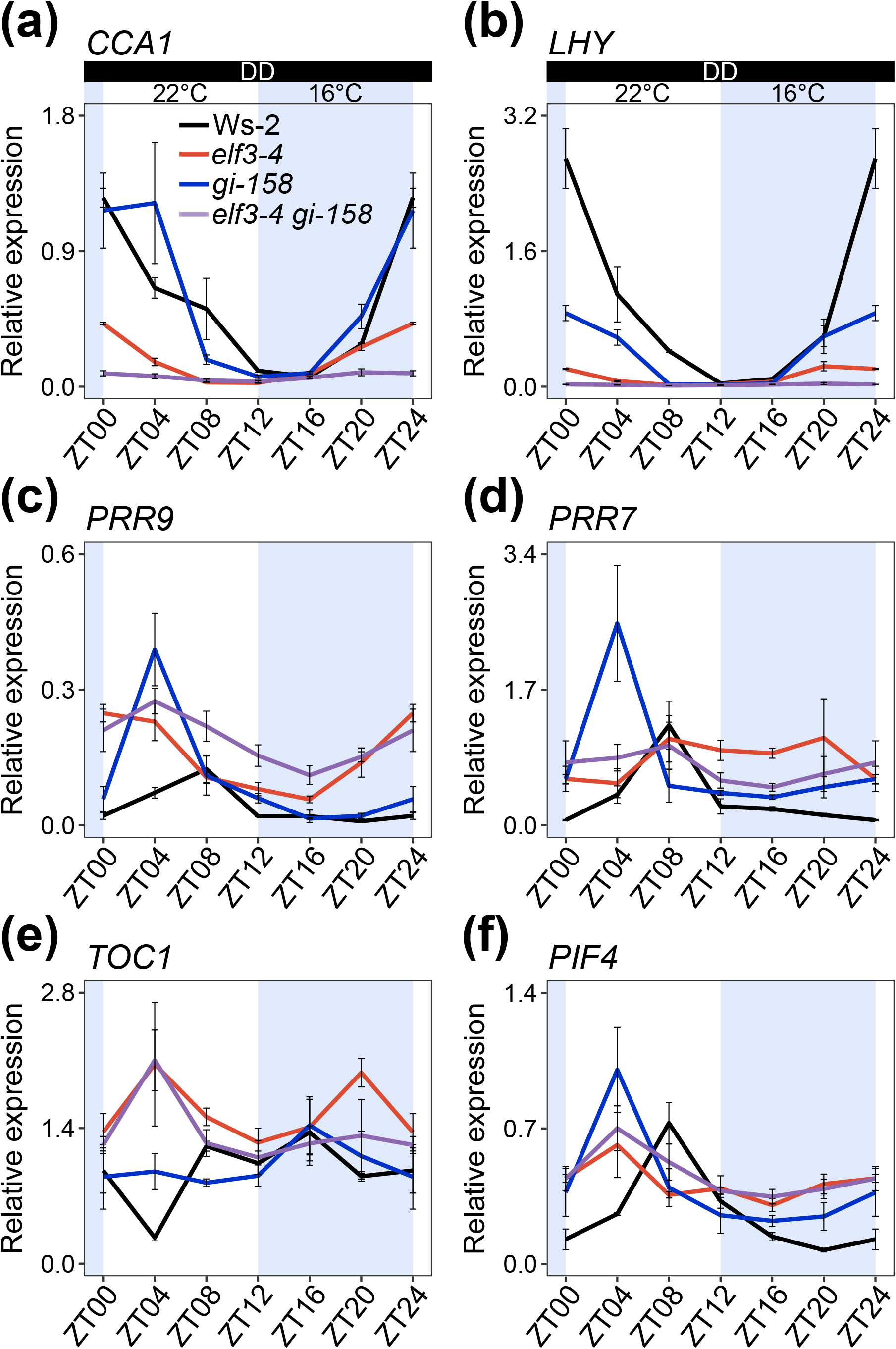
*ELF3* is required for oscillator’s responsiveness to temperature changes in darkness. (a-f) Transcript dynamics of key clock oscillator genes *CCA1* (a), *LHY* (b), *PRR9* (c), *PRR7* (d), *TOC1* (e) and major growth promoter *PIF4* (f). Ws-2, *elf3-4*, *gi-158* and *elf3-4 gi-158* seedlings were grown in DD under thermocycles for 8 d. On day 9, starting from ZT00, samples were harvested every 4 h. Non-shaded areas represent warm period (22°C), whereas blue-shaded areas represent cold period (16°C). Expression levels were normalized to *PP2A*. Error bars indicate SEM (n = 3) of three biological replicates.

**Fig. S5.**
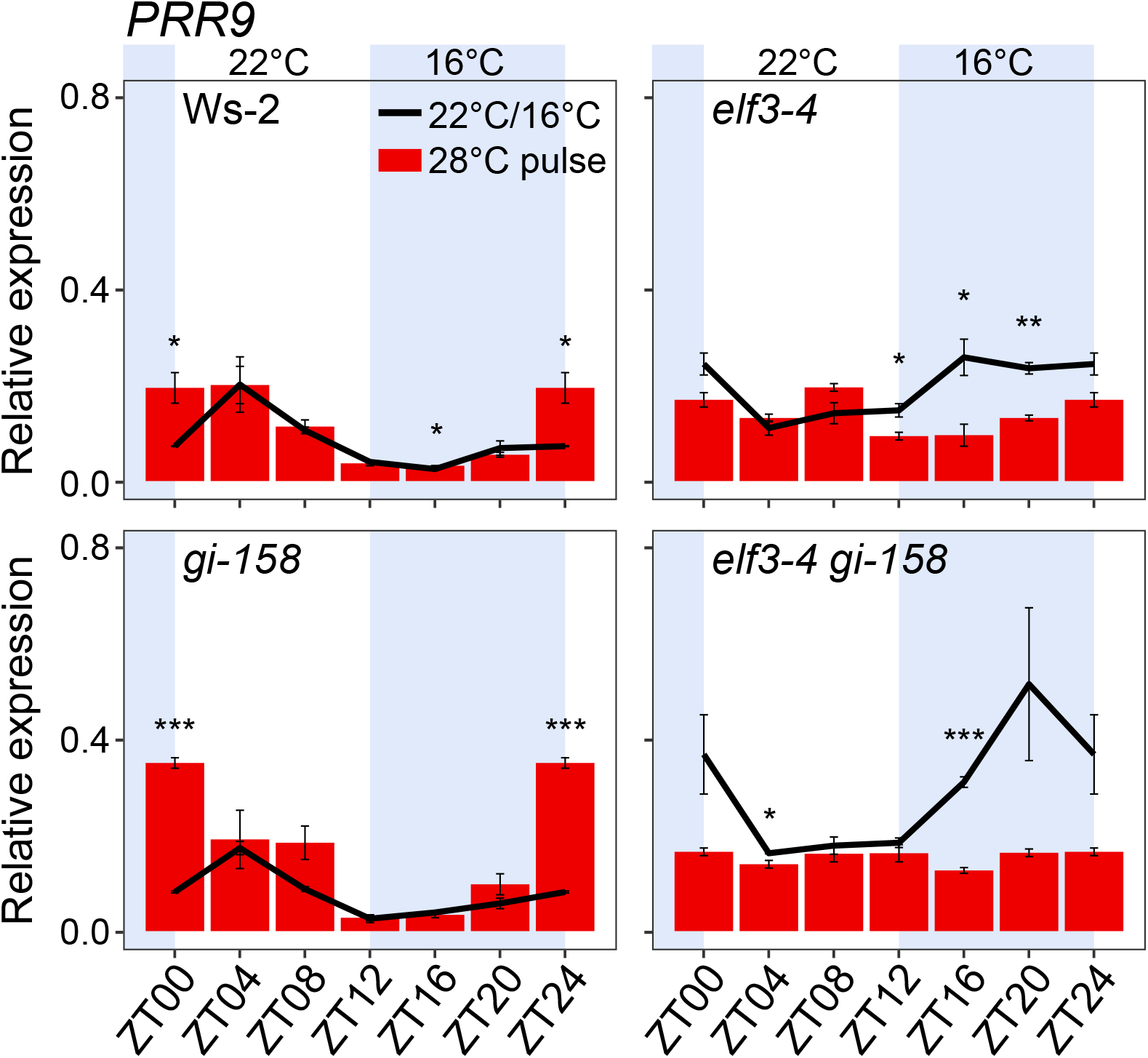
*ELF3* is essential for precise gating of the temperature signals under thermocycles. Effect of the temperature pulse at the specified ZTs on the expression of *PRR9*. Ws-2, *elf3-4*, *gi-158* and *elf3-4 gi-158* seedlings were grown in LL under thermocycles for 8 d. On day 9, the seedlings were either treated with a 4 h temperature pulse (28°C pulse) at indicated ZTs or were kept under the same conditions (no treatment, 22°C/16°C) before samples were harvested. At indicated ZTs, red bars represent gene expression levels after treatment with a temperature pulse, whereas black lines represent gene expression levels at the same time without treatment. Non-shaded areas represent warm period (22°C), whereas blue-shaded areas represent cold period (16°C). Expression levels were normalized to *PP2A*. Error bars indicate SEM (n = 3) of three biological replicates. Asterisks above lines or bars indicate significant differences (*, *P* < 0.05; **, *P* < 0.01; ***, *P* < 0.001; Student’s *t*-test).

**Fig. S6.**
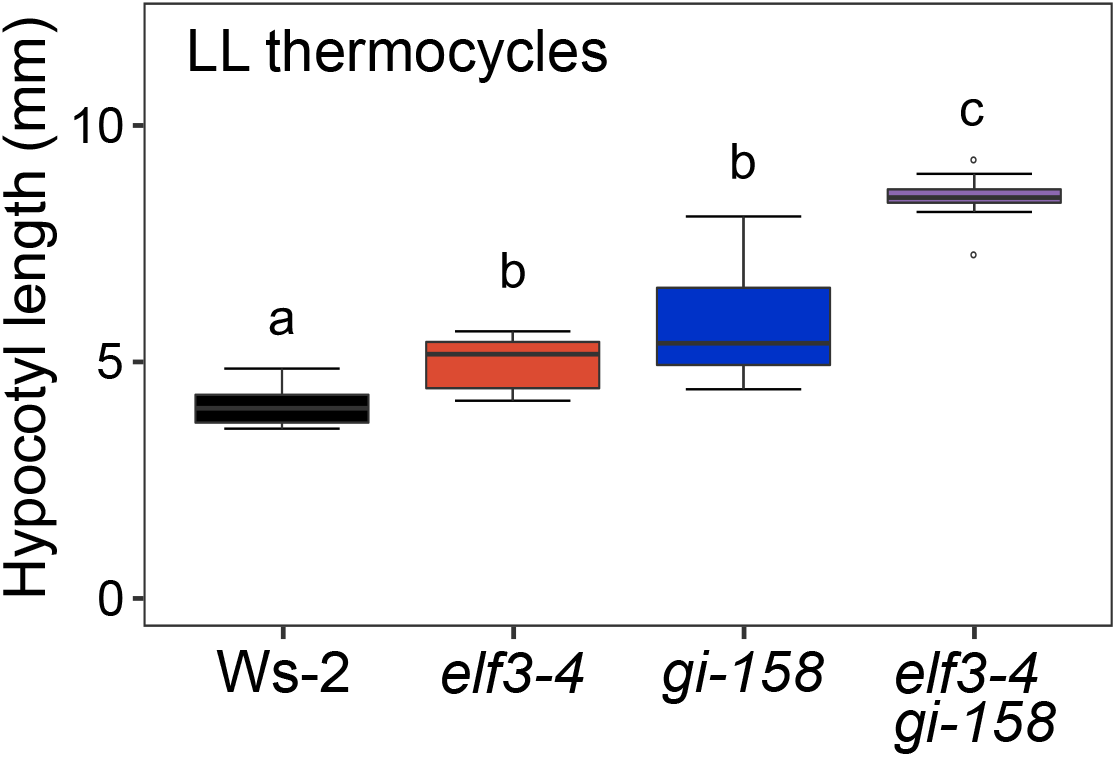
*ELF3* and *GI* display additive effect under thermocycles in constant light. Quantification of hypocotyl length of 6-d-old Ws-2, *elf3-4*, *gi-158* and *elf3-4 gi-158* seedlings grown in LL (30 μmol m^−2^s^−1^), under 12 h 22°C : 12 h 16°C thermocycles. Box plots show medians and interquartile ranges. Outliers (greater than 1.5x interquartile range) are marked with open circles. Different letters above the boxes indicate significant differences (one-way ANOVA with Tukey’s HSD test, *P* < 0.05).

**Table S1.** Rhythmic growth under thermocycles requires *ELF3*.

| Time (h)/ZT* | 48/ZT00 | 52/ZT04 | 56/ZT08 | 60/ZT12 | 64/ZT16 | 68/ZT20 | 72/ZT24 |  |
| --- | --- | --- | --- | --- | --- | --- | --- | --- |
| Genotype | Growth (mm h<sup>-1</sup>) ± SEM (n = 8) |  |  |  |  |  |  | *P* |
| Ws-2 | 0.0075 ± 0.0056 a<sup>†</sup> | 0.0533 ± 0.0250 ab | **0.0996 ± 0.0353** b | 0.0584 ± 0.0199 ab | 0.0164 ± 0.0092 a | 0.0100 ± 0.0071 a | 0.0042 ± 0.0034 a | 0.004 |
| *elf3-4* | 0.0273 ± 0.0070 | 0.0448 ± 0.0113 | 0.0469 ± 0.0071 | 0.0432 ± 0.0113 | 0.0258 ± 0.0058 | 0.0244 ± 0.0074 | 0.0278 ± 0.0114 | 0.299 |
| *gi-158* | 0.0082 ± 0.0065 a | 0.0795 ± 0.0125 c | 0.0746 ± 0.0192 bc | 0.0533 ± 0.0178 abc | 0.0318 ± 0.0078 abc | 0.0126 ± 0.0072 ab | 0.0264 ± 0.0206 abc | 0.002 |
| *elf3-4 gi-158* | 0.0788 ± 0.0202 | 0.0858 ± 0.0148 | 0.0971 ± 0.0112 | 0.0806 ± 0.0150 | 0.0595 ± 0.0111 | 0.0476 ± 0.0144 | 0.0656 ± 0.0174 | 0.300 |
|  | 72/ZT00 | 76/ZT04 | 80/ZT08 | 84/ZT12 | 88/ZT16 | 92/ZT20 | 96/ZT24 |  |
| Ws-2 | 0.0042 ± 0.0034 a | 0.0361 ± 0.0138 ab | **0.0836 ± 0.0187** b | 0.0410 ± 0.0208 ab | 0.0095 ± 0.0055 a | 0.0128 ± 0.0128 a | 0.0019 ± 0.0011 a | 0.003 |
| *elf3-4* | 0.0278 ± 0.0114 | 0.0239 ± 0.0086 | 0.0258 ± 0.0081 | 0.0150 ± 0.0064 | 0.0364 ± 0.0144 | 0.0382 ± 0.0108 | 0.0205 ± 0.0073 | 0.649 |
| *gi-158* | 0.0264 ± 0.0206 a | 0.0589 ± 0.0199 ab | **0.0941 ± 0.0100** b | 0.0540 ± 0.0162 ab | 0.0309 ± 0.0123 a | 0.0196 ± 0.0098 a | 0.0062 ± 0.047 a | 0.002 |
| *elf3-4 gi-158* | 0.0656 ± 0.0174 | 0.0641 ± 0.0115 | 0.0756 ± 0.0145 | 0.0725 ± 0.0183 | 0.0529 ± 0.0163 | 0.0575 ± 0.0129 | 0.0296 ± 0.0127 | 0.399 |
|  | 96/ZT00 | 100/ZT04 | 104/ZT08 | 108/ZT12 | 112/ZT16 | 116/ZT20 | 120/ZT24 |  |
| Ws-2 | 0.0019 ± 0.0011 a | 0.0408 ± 0.0121 ab | 0.0405 ± 0.0120 ab | **0.0711 ± 0.0231** b | 0.0049 ± 0.0043 a | 0.0058 ± 0.0056 a | 0.0239 ± 0.0131 ab | 0.001 |
| *elf3-4* | 0.0205 ± 0.0073 | 0.0314 ± 0.0119 | 0.0276 ± 0.0070 | 0.0255 ± 0.0064 | 0.0425 ± 0.0125 | 0.0168 ± 0.0064 | 0.0108 ± 0.0035 | 0.193 |
| *gi-158* | 0.0062 ± 0.0047 | 0.0461 ± 0.0117 | 0.0535 ± 0.0190 | 0.0530 ± 0.0149 | 0.0155 ± 0.0073 | 0.0223 ± 0.0098 | 0.0164 ± 0.0113 | 0.024 |
| *elf3-4 gi-158* | 0.0296 ± 0.0127 ab | 0.0804 ± 0.0102 b | 0.0776 ± 0.0216 ab | 0.0401 ± 0.0079 | 0.0530 ± 0.0147 ab | 0.0356 ± 0.0129 ab | 0.0211 ± 0.0107 a | 0.017 |
|  | 120/ZT00 | 124/ZT04 | 128/ZT08 | 132/ZT12 | 136/ZT16 | 140/ZT20 | 144/ZT24 |  |
| Ws-2 | 0.0239 ± 0.0131 a | 0.0296 ± 0.0154 ab | **0.0668 ± 0.0116** b | 0.0366 ± 0.0078 ab | 0.0019 ± 0.0013 a | 0.0064 ± 0.0039 a | 0.0009 ± 0.0005 a | 0.000 |
| *elf3-4* | 0.0108 ± 0.0035 | 0.0295 ± 0.0120 | 0.0334 ± 0.0105 | 0.0172 ± 0.0059 | 0.0162 ± 0.0056 | 0.0204 ± 0.0090 | 0.0142 ± 0.0090 | 0.458 |
| *gi-158* | 0.0164 ± 0.0113 ab | 0.0534 ± 0.0124 bc | **0.0861 ± 0.0219** c | 0.0335 ± 0.0119 ab | 0.0076 ± 0.0050 ab | 0.0000 ± 0.0000 a | 0.0000 ± 0.0000 a | 0.000 |
| *elf3-4 gi-158* | 0.0211 ± 0.0107 | 0.0394 ± 0.0109 | 0.0428 ± 0.0125 | 0.0193 ± 0.0070 | 0.0268 ± 0.0088 | 0.0257 ± 0.0112 | 0.0032 ± 0.0032 | 0.104 |

\*Seedlings were grown under 12 h 22°C: 12 h 16°C thermocycles in constant light. Indicated time points (every 4 h) were selected for statistics.
<sup>†</sup>Different letters indicate significant differences in growth rate within indicated time points per genotype per day (one-way ANOVA with Tukey's HSD test).
The value with the highest mean, significantly higher than the values of more than three other time points, is considered as a peak (shown in bold).

**Table S2.** *ELF3* is required for oscillator’s responsiveness to temperature change in constant light.

| ZT* | ZT00/24 | ZT04 | ZT08 | ZT12 | ZT16 | ZT20 |  |
| --- | --- | --- | --- | --- | --- | --- | --- |
| Genotype | CCA1 relative expression ± SEM (n = 3) | | | | | | *P* |
| Ws-2 | **2.2428 ± 0.7727** a<sup>†</sup> | 1.5875 ± 0.1476 ab | 0.2306 ± 0.0583 b | 0.0621 ± 0.0174 b | 0.1062 ± 0.0244 b | 0.3102 ± 0.0613 b | 0.001 |
| *elf3-4* | 0.1007 ± 0.0175 | 0.0976 ± 0.0016 | 0.0719 ± 0.0139 | 0.0610 ± 0.0051 | 0.0677 ± 0.0063 | 0.1059 ± 0.0165 | 0.064 |
| *gi-158* | **1.2499 ± 0.1559** a | 0.7480 ± 0.1260 b | 0.0544 ± 0.0090 c | 0.0133 ± 0.0013 c | 0.0565 ± 0.0098 c | 0.5993 ± 0.1529 b | 0.000 |
| *elf3-4 gi-158* | 0.0667 ± 0.0070 a | 0.0528 ± 0.0008 a | 0.0449 ± 0.0023 a | 0.0542 ± 0.0034 a | 0.0780 ± 0.0062 ab | **0.1194 ± 0.0202** b | 0.001 |
| | *LHY* relative expression ± SEM (n = 3) | | | | | | |
| Ws-2 | **4.4052 ± 0.9873** a | 1.1700 ± 0.1084 b | 0.5359 ± 0.2465 b | 0.0787 ± 0.0286 b | 0.1578 ± 0.0230 b | 0.9713 ± 0.1112 b | 0.000 |
| *elf3-4* | 0.0854 ± 0.0059 a | 0.1132 ± 0.0343 ab | 0.0840 ± 0.0175 a | 0.0521 ± 0.0139 a | 0.0700 ± 0.0029 a | **0.1930 ± 0.0287** b | 0.003 |
| *gi-158* | **2.9962 ± 1.4138** a | 0.2764 ± 0.0165 ab | 0.0689 ± 0.0215 b | 0.0083 ± 0.0000 b | 0.0746 ± 0.0132 b | 1.5321 ± 0.1575 ab | 0.017 |
| *elf3-4 gi-158* | 0.0110 ± 0.0004 ab | 0.0116 ± 0.0017 ab | 0.0040 ± 0.0004 b | 0.0088 ± 0.0040 ab | 0.0090 ± 0.0004 ab | 0.0169 ± 0.0012 a | 0.002 |
| | *PRR9* relative expression ± SEM (n = 3) | | | | | | |
| Ws-2 | 0.0328 ± 0.0126 ab | **0.2447 ± 0.0509** c | 0.1253 ± 0.0025 b | 0.0539 ± 0.0156 ab | 0.0212 ± 0.0051 ab | 0.0177 ± 0.0022 a | 0.000 |
| *elf3-4* | 0.1354 ± 0.0079 | 0.1033 ± 0.0056 | 0.1221 ± 0.0079 | 0.1241 ± 0.0137 | 0.1445 ± 0.0414 | 0.1509 ± 0.0163 | 0.665 |
| *gi-158* | 0.0410 ± 0.0049 a | **0.1950 ± 0.0329** c | 0.1186 ± 0.0087 b | 0.0264 ± 0.0051 a | 0.0253 ± 0.0008 a | 0.0427 ± 0.0071 a | 0.000 |
| *elf3-4 gi-158* | 0.1948 ± 0.0350 | 0.1635 ± 0.0275 | 0.1483 ± 0.0124 | 0.2593 ± 0.0581 | 0.2118 ± 0.0265 | 0.3008 ± 0.0359 | 0.066 |
| | *PRR7* relative expression ± SEM (n = 3) | | | | | | |
| Ws-2 | 0.0463 ± 0.0070 ab | 0.2812 ± 0.0129 c | **0.5575 ± 0.0078** d | 0.2657 ± 0.0544 c | 0.1247 ± 0.0446 b | 0.0398 ± 0.0060 a | 0.000 |
| *elf3-4* | 0.3115 ± 0.0218 ab | 0.3402 ± 0.0641 ab | 0.4039 ± 0.0239 ab | 0.2385 ± 0.0488 a | 0.3555 ± 0.0302 ab | 0.4791 ± 0.0552 b | 0.034 |
| *gi-158* | 0.0721 ± 0.0275 a | 0.4797 ± 0.1555 ab | **0.9709 ± 0.3374** b | 0.1570 ± 0.0254 a | 0.0402 ± 0.0104 a | 0.0418 ± 0.0038 a | 0.001 |
| *elf3-4 gi-158* | 0.4483 ± 0.0280 ab | 0.5457 ± 0.1809 ab | 0.2909 ± 0.0169 a | 0.2832 ± 0.0463 a | 0.3623 ± 0.0222 ab | 0.7201 ± 0.0777 b | 0.023 |

**Table S2. (continued)**
| ZT | ZT00/24 | ZT04 | ZT08 | ZT12 | ZT16 | ZT20 |  |
| --- | --- | --- | --- | --- | --- | --- | --- |
| Genotype | *TOC1* relative expression ± SEM (n = 3) | | | | | | *P* |
| Ws-2 | 0.2311 ± 0.0307 ab | 0.1547 ± 0.0156 a | 0.2169 ± 0.0225 ab | 0.4489 ± 0.0443 bc | **0.5571 ± 0.1131** c | 0.2739 ± 0.0244 ab | 0.001 |
| *elf3-4* | 0.3609 ± 0.0158 | 0.2800 ± 0.0134 | 0.4263 ± 0.0825 | 0.3676 ± 0.0445 | 0.4075 ± 0.0973 | 0.3496 ± 0.0056 | 0.675 |
| *gi-158* | 0.1724 ± 0.0089 ab | 0.1092 ± 0.0148 a | 0.2894 ± 0.0263 bc | **0.4294 ± 0.0481** d | 0.3179 ± 0.0350 cd | 0.2090 ± 0.0186 abc | 0.000 |
| *elf3-4 gi-158* | 0.3555 ± 0.0100 ab | 0.2486 ± 0.0194 a | 0.3904 ± 0.0263 b | 0.2658 ± 0.0463 ab | 0.2906 ± 0.0285 ab | 0.3292 ± 0.0236 ab | 0.027 |
| | *PIF4* relative expression ± SEM (n = 3) | | | | | | |
| Ws-2 | 0.0286 ± 0.0068 a | 0.2991 ± 0.0551 b | **0.4161 ± 0.0241** b | 0.2523 ± 0.0563 b | 0.0797 ± 0.0186 a | 0.0217 ± 0.0062 a | 0.000 |
| *elf3-4* | 0.2105 ± 0.0184 | 0.2620 ± 0.0064 | 0.2830 ± 0.0411 | 0.3101 ± 0.0306 | 0.2288 ± 0.0269 | 0.2284 ± 0.0180 | 0.129 |
| *gi-158* | 0.0482 ± 0.0059 ab | 0.4175 ± 0.0546 c | **0.6012 ± 0.0616** d | 0.2010 ± 0.0291 b | 0.0333 ± 0.0041 ab | 0.0177 ± 0.0028 a | 0.000 |
| *elf3-4 gi-158* | 0.3739 ± 0.0430 | 0.3722 ± 0.0271 | 0.3904 ± 0.0172 | 0.4293 ± 0.0407 | 0.3366 ± 0.0750 | 0.4019 ± 0.0131 | 0.734 |

\*Seedlings were grown under 12 h 22°C: 12 h 16°C thermocycles in constant light for 8 d. On day 9, starting from ZT00, samples were harvested every 4 h. Expression levels were normalized to *PP2A*.
†Different letters indicate significant differences in expression level within indicated time points per genotype (one-way ANOVA with Tukey's HSD test). The value with the highest mean, significantly higher than the values of more than three other time points, is considered as a peak (shown in bold).

**Table S3.**
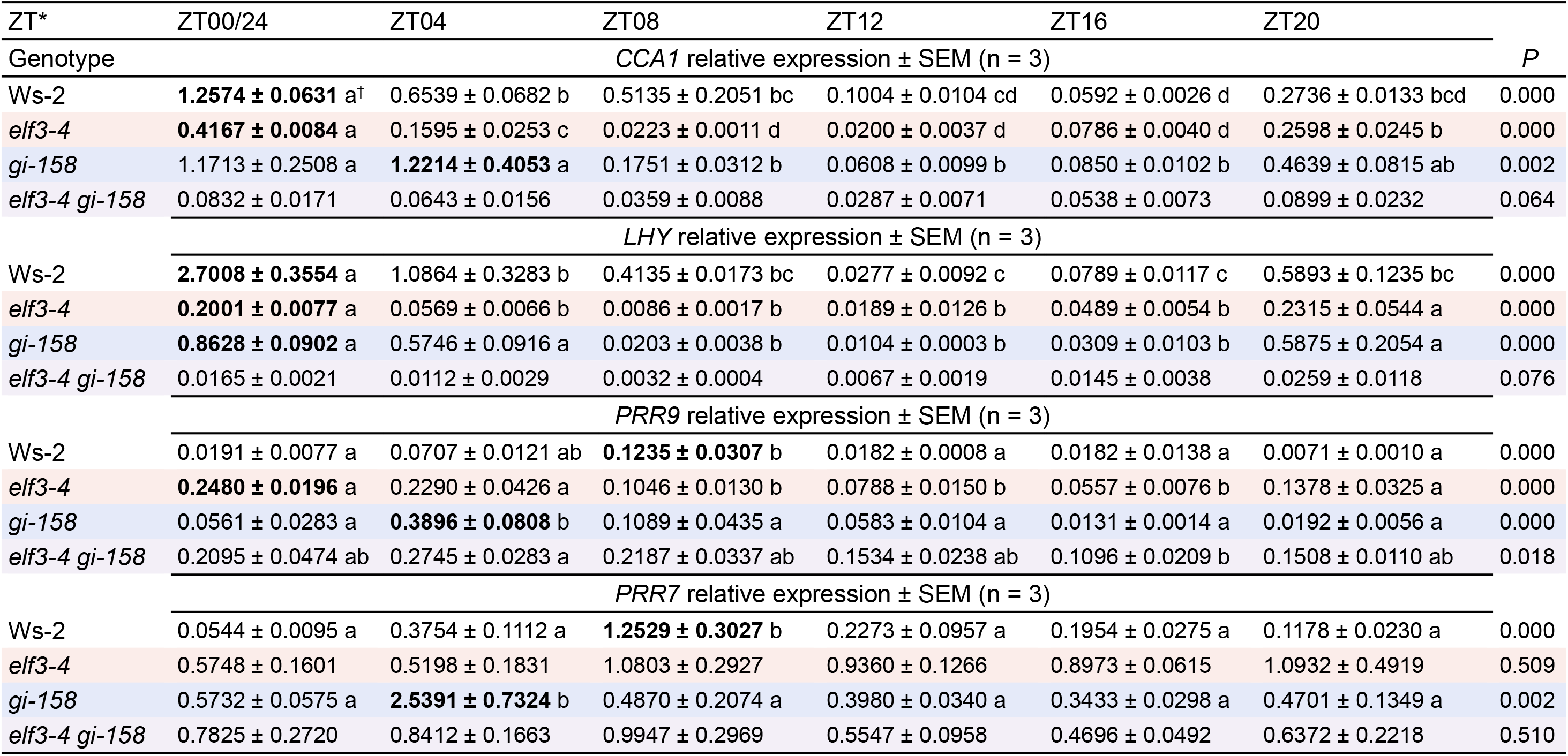
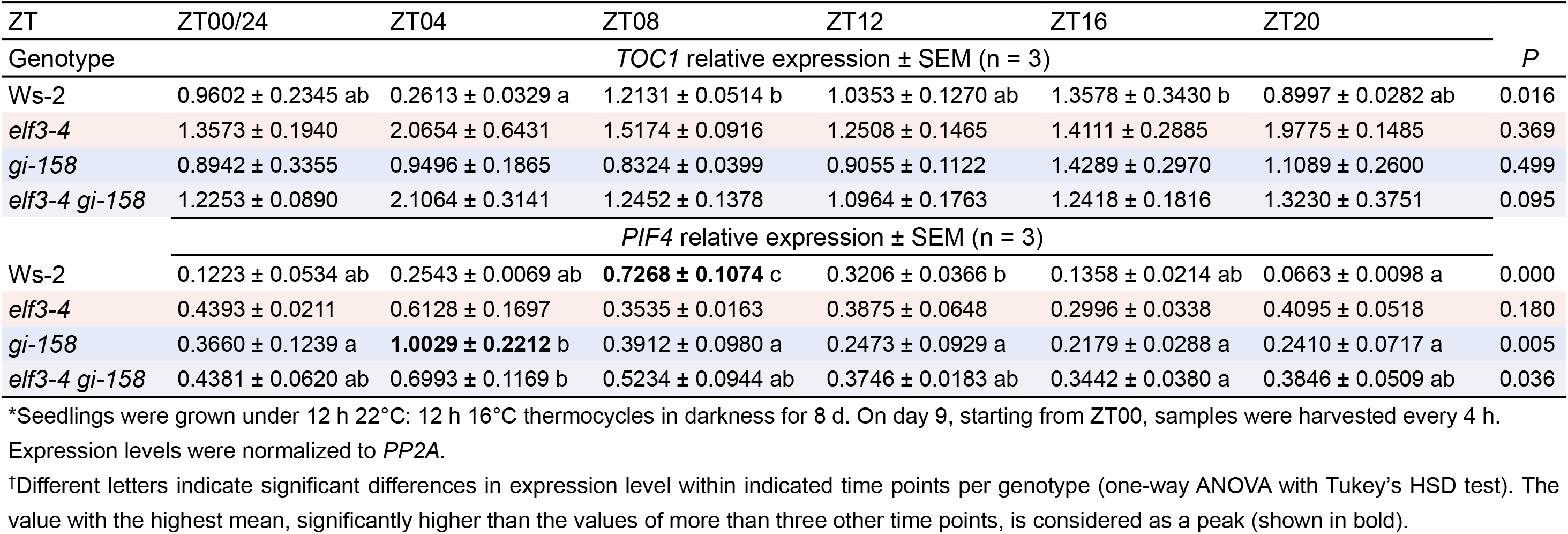
*ELF3* is required for oscillator’s responsiveness to temperature change in darkness.

| ZT* | ZT00/24 | ZT04 | ZT08 | ZT12 | ZT16 | ZT20 |  |
| --- | --- | --- | --- | --- | --- | --- | --- |
| Genotype | CCA1 relative expression ± SEM (n = 3) | | | | | | P |
| Ws-2 | **1.2574 ± 0.0631** a<sup>†</sup> | 0.6539 ± 0.0682 b | 0.5135 ± 0.2051 bc | 0.1004 ± 0.0104 cd | 0.0592 ± 0.0026 d | 0.2736 ± 0.0133 bcd | 0.000 |
| *elf3-4* | **0.4167 ± 0.0084** a | 0.1595 ± 0.0253 c | 0.0223 ± 0.0011 d | 0.0200 ± 0.0037 d | 0.0786 ± 0.0040 d | 0.2598 ± 0.0245 b | 0.000 |
| *gi-158* | 1.1713 ± 0.2508 a | **1.2214 ± 0.4053** a | 0.1751 ± 0.0312 b | 0.0608 ± 0.0099 b | 0.0850 ± 0.0102 b | 0.4639 ± 0.0815 ab | 0.002 |
| *elf3-4 gi-158* | 0.0832 ± 0.0171 | 0.0643 ± 0.0156 | 0.0359 ± 0.0088 | 0.0287 ± 0.0071 | 0.0538 ± 0.0073 | 0.0899 ± 0.0232 | 0.064 |
| | LHY relative expression ± SEM (n = 3) | | | | | | |
| Ws-2 | **2.7008 ± 0.3554** a | 1.0864 ± 0.3283 b | 0.4135 ± 0.0173 bc | 0.0277 ± 0.0092 c | 0.0789 ± 0.0117 c | 0.5893 ± 0.1235 bc | 0.000 |
| *elf3-4* | **0.2001 ± 0.0077** a | 0.0569 ± 0.0066 b | 0.0086 ± 0.0017 b | 0.0189 ± 0.0126 b | 0.0489 ± 0.0054 b | 0.2315 ± 0.0544 a | 0.000 |
| *gi-158* | **0.8628 ± 0.0902** a | 0.5746 ± 0.0916 a | 0.0203 ± 0.0038 b | 0.0104 ± 0.0003 b | 0.0309 ± 0.0103 b | 0.5875 ± 0.2054 a | 0.000 |
| *elf3-4 gi-158* | 0.0165 ± 0.0021 | 0.0112 ± 0.0029 | 0.0032 ± 0.0004 | 0.0067 ± 0.0019 | 0.0145 ± 0.0038 | 0.0259 ± 0.0118 | 0.076 |
| | PRR9 relative expression ± SEM (n = 3) | | | | | | |
| Ws-2 | 0.0191 ± 0.0077 a | 0.0707 ± 0.0121 ab | **0.1235 ± 0.0307** b | 0.0182 ± 0.0008 a | 0.0182 ± 0.0138 a | 0.0071 ± 0.0010 a | 0.000 |
| *elf3-4* | **0.2480 ± 0.0196** a | 0.2290 ± 0.0426 a | 0.1046 ± 0.0130 b | 0.0788 ± 0.0150 b | 0.0557 ± 0.0076 b | 0.1378 ± 0.0325 a | 0.000 |
| *gi-158* | 0.0561 ± 0.0283 a | **0.3896 ± 0.0808** b | 0.1089 ± 0.0435 a | 0.0583 ± 0.0104 a | 0.0131 ± 0.0014 a | 0.0192 ± 0.0056 a | 0.000 |
| *elf3-4 gi-158* | 0.2095 ± 0.0474 ab | 0.2745 ± 0.0283 a | 0.2187 ± 0.0337 ab | 0.1534 ± 0.0238 ab | 0.1096 ± 0.0209 b | 0.1508 ± 0.0110 ab | 0.018 |
| | PRR7 relative expression ± SEM (n = 3) | | | | | | |
| Ws-2 | 0.0544 ± 0.0095 a | 0.3754 ± 0.1112 a | **1.2529 ± 0.3027** b | 0.2273 ± 0.0957 a | 0.1954 ± 0.0275 a | 0.1178 ± 0.0230 a | 0.000 |
| *elf3-4* | 0.5748 ± 0.1601 | 0.5198 ± 0.1831 | 1.0803 ± 0.2927 | 0.9360 ± 0.1266 | 0.8973 ± 0.0615 | 1.0932 ± 0.4919 | 0.509 |
| *gi-158* | 0.5732 ± 0.0575 a | **2.5391 ± 0.7324** b | 0.4870 ± 0.2074 a | 0.3980 ± 0.0340 a | 0.3433 ± 0.0298 a | 0.4701 ± 0.1349 a | 0.002 |
| *elf3-4 gi-158* | 0.7825 ± 0.2720 | 0.8412 ± 0.1663 | 0.9947 ± 0.2969 | 0.5547 ± 0.0958 | 0.4696 ± 0.0492 | 0.6372 ± 0.2218 | 0.510 |

**Table S3. (continued)**
| ZT | ZT00/24 | ZT04 | ZT08 | ZT12 | ZT16 | ZT20 |  |
| --- | --- | --- | --- | --- | --- | --- | --- |
| Genotype | *TOC1* relative expression ± SEM (n = 3) | | | | | | *P* |
| Ws-2 | 0.9602 ± 0.2345 ab | 0.2613 ± 0.0329 a | 1.2131 ± 0.0514 b | 1.0353 ± 0.1270 ab | 1.3578 ± 0.3430 b | 0.8997 ± 0.0282 ab | 0.016 |
| *elf3-4* | 1.3573 ± 0.1940 | 2.0654 ± 0.6431 | 1.5174 ± 0.0916 | 1.2508 ± 0.1465 | 1.4111 ± 0.2885 | 1.9775 ± 0.1485 | 0.369 |
| *gi-158* | 0.8942 ± 0.3355 | 0.9496 ± 0.1865 | 0.8324 ± 0.0399 | 0.9055 ± 0.1122 | 1.4289 ± 0.2970 | 1.1089 ± 0.2600 | 0.499 |
| *elf3-4 gi-158* | 1.2253 ± 0.0890 | 2.1064 ± 0.3141 | 1.2452 ± 0.1378 | 1.0964 ± 0.1763 | 1.2418 ± 0.1816 | 1.3230 ± 0.3751 | 0.095 |
| | *PIF4* relative expression ± SEM (n = 3) | | | | | | |
| Ws-2 | 0.1223 ± 0.0534 ab | 0.2543 ± 0.0069 ab | **0.7268 ± 0.1074** c | 0.3206 ± 0.0366 b | 0.1358 ± 0.0214 ab | 0.0663 ± 0.0098 a | 0.000 |
| *elf3-4* | 0.4393 ± 0.0211 | 0.6128 ± 0.1697 | 0.3535 ± 0.0163 | 0.3875 ± 0.0648 | 0.2996 ± 0.0338 | 0.4095 ± 0.0518 | 0.180 |
| *gi-158* | 0.3660 ± 0.1239 a | **1.0029 ± 0.2212** b | 0.3912 ± 0.0980 a | 0.2473 ± 0.0929 a | 0.2179 ± 0.0288 a | 0.2410 ± 0.0717 a | 0.005 |
| *elf3-4 gi-158* | 0.4381 ± 0.0620 ab | 0.6993 ± 0.1169 b | 0.5234 ± 0.0944 ab | 0.3746 ± 0.0183 ab | 0.3442 ± 0.0380 a | 0.3846 ± 0.0509 ab | 0.036 |

\*Seedlings were grown under 12 h 22°C: 12 h 16°C thermocycles in darkness for 8 d. On day 9, starting from ZT00, samples were harvested every 4 h.
Expression levels were normalized to *PP2A*.
†Different letters indicate significant differences in expression level within indicated time points per genotype (one-way ANOVA with Tukey's HSD test). The value with the highest mean, significantly higher than the values of more than three other time points, is considered as a peak (shown in bold).

## References

Adams S, Manfield I, Stockley P, Carré IA. 2015. Revised Morning Loops of the Arabidopsis Circadian Clock Based on Analyses of Direct Regulatory Interactions. PLOS ONE 10(12): e0143943.

Alabadı́ D, Oyama T, Yanovsky MJ, Harmon FG, Más P, Kay SA. 2001. Reciprocal Regulation Between TOC1 and LHY/CCA1 Within the Arabidopsis Circadian Clock. Science 293(5531): 880–883.

Anwer MU, Boikoglou E, Herrero E, Hallstein M, Davis AM, Velikkakam James G, Nagy F, Davis SJ, Mockler TC. 2014. Natural variation reveals that intracellular distribution of ELF3 protein is associated with function in the circadian clock. eLife.

Anwer MU, Davis A, Davis SJ, Quint M. 2020. Photoperiod sensing of the circadian clock is controlled by EARLY FLOWERING 3 and GIGANTEA. The Plant Journal 101(6): 1397–1410.

Avello PA, Davis SJ, Ronald J, Pitchford JW. 2019. Heat the Clock: Entrainment and Compensation in Arabidopsis Circadian Rhythms. Journal of circadian rhythms 17: 5–5.

Bendix C, Marshall Carine M, Harmon Frank G. 2015. Circadian Clock Genes Universally Control Key Agricultural Traits. Molecular Plant 8(8): 1135–1152.

Box Mathew S, Huang BE, Domijan M, Jaeger Katja E, Khattak Asif K, Yoo Seong J, Sedivy Emma L, Jones DM, Hearn Timothy J, Webb Alex AR, et al. 2015. ELF3 Controls Thermoresponsive Growth in Arabidopsis. Current Biology(0).

Covington MF, Maloof JN, Straume M, Kay SA, Harmer SL. 2008. Global transcriptome analysis reveals circadian regulation of key pathways in plant growth and development. Genome Biology 9(8): R130.

Delker C, van Zanten M, Quint M. 2017. Thermosensing enlightened. Trends in Plant Science 22(3): 185–187.

Eckardt NA 2005. Temperature entrainment of the Arabidopsis circadian clock: Am Soc Plant Biol.

Ezer D, Jung J-H, Lan H, Biswas S, Gregoire L, Box MS, Charoensawan V, Cortijo S, Lai X, Stöckle D, et al. 2017. The evening complex coordinates environmental and endogenous signals in Arabidopsis. 3: 17087.

Faure S, Turner AS, Gruszka D, Christodoulou V, Davis SJ, von Korff M, Laurie DA. 2012. Mutation at the circadian clock gene EARLY MATURITY 8 adapts domesticated barley (Hordeum vulgare) to short growing seasons. Proceedings of the National Academy of Sciences.

Fowler S, Lee K, Onouchi H, Samach A, Richardson K, Morris B, Coupland G, Putterill J. 1999. GIGANTEA: a circadian clock-controlled gene that regulates photoperiodic flowering in Arabidopsis and encodes a protein with several possible membrane-spanning domains. EMBO J 18(17): 4679–4688.

Gil KE, Park CM. 2019. Thermal adaptation and plasticity of the plant circadian clock. New Phytologist 221(3): 1215–1229.

Herrero E, Kolmos E, Bujdoso N, Yuan Y, Wang M, Berns MC, Uhlworm H, Coupland G, Saini R, Jaskolski M, et al. 2012. EARLY FLOWERING4 Recruitment of EARLY FLOWERING3 in the Nucleus Sustains the Arabidopsis Circadian Clock. The Plant Cell Online.

Hicks KA, Albertson TM, Wagner DR. 2001. EARLY FLOWERING3 encodes a novel protein that regulates circadian clock function and flowering in Arabidopsis. The Plant Cell 13(6): 1281–1292.

Huang H, Nusinow DA. 2016. Into the Evening: Complex Interactions in the Arabidopsis Circadian Clock. Trends in Genetics 32(10): 674–686.

Huang W, Perez-Garcia P, Pokhilko A, Millar AJ, Antoshechkin I, Riechmann JL, Mas P. 2012. Mapping the Core of the Arabidopsis Circadian Clock Defines the Network Structure of the Oscillator. Science 336(6077): 75–79.

Huq E, Tepperman JM, Quail PH. 2000. GIGANTEA is a nuclear protein involved in phytochrome signaling in Arabidopsis. Proc Natl Acad Sci U S A 97(17): 9789–9794.

Jung J-H, Barbosa AD, Hutin S, Kumita JR, Gao M, Derwort D, Silva CS, Lai X, Pierre E, Geng F, et al. 2020. A prion-like domain in ELF3 functions as a thermosensor in Arabidopsis. Nature.

Jung J-H, Domijan M, Klose C, Biswas S, Ezer D, Gao M, Khattak AK, Box MS, Charoensawan V, Cortijo S, et al. 2016. Phytochromes function as thermosensors in Arabidopsis. Science.

Kamioka M, Takao S, Suzuki T, Taki K, Higashiyama T, Kinoshita T, Nakamichi N. 2016. Direct Repression of Evening Genes by CIRCADIAN CLOCK-ASSOCIATED1 in the Arabidopsis Circadian Clock. The Plant Cell 28(3): 696.

Kim W-Y, Fujiwara S, Suh S-S, Kim J, Kim Y, Han L, David K, Putterill J, Nam HG, Somers DE. 2007. ZEITLUPE is a circadian photoreceptor stabilized by GIGANTEA in blue light. Nature 449(7160): 356–360.

Kim Y, Lim J, Yeom M, Kim H, Kim J, Wang L, Kim Woe Y, Somers David E, Nam Hong G. 2013. ELF4 Regulates GIGANTEA Chromatin Access through Subnuclear Sequestration. Cell Reports 3(3): 671–677.

Kim Y, Yeom M, Kim H, Lim J, Koo HJ, Hwang D, Somers D, Nam HG. 2012. GIGANTEA and EARLY FLOWERING 4 in Arabidopsis Exhibit Differential Phase-Specific Genetic Influences over a Diurnal Cycle. Molecular Plant.

Kolmos E, Herrero E, Bujdoso N, Millar AJ, Toth R, Gyula P, Nagy F, Davis SJ. 2011. A Reduced-Function Allele Reveals That EARLY FLOWERING3 Repressive Action on the Circadian Clock Is Modulated by Phytochrome Signals in Arabidopsis. Plant Cell.

Legris M, Klose C, Burgie ES, Rojas CCR, Neme M, Hiltbrunner A, Wigge PA, Schäfer E, Vierstra RD, Casal JJ. 2016. Phytochrome B integrates light and temperature signals in Arabidopsis. Science 354(6314): 897–900.

Lincoln C, Britton JH, Estelle M. 1990. Growth and development of the axr1 mutants of Arabidopsis. The Plant Cell 2(11): 1071–1080.

Lippmann R, Babben S, Menger A, Delker C, Quint M. 2019. Development of Wild and Cultivated Plants under Global Warming Conditions. Current Biology 29(24): R1326–R1338.

Liu XL, Covington MF, Fankhauser C, Chory J, Wagner DR. 2001. ELF3 encodes a circadian clock–regulated nuclear protein that functions in an Arabidopsis PHYB signal transduction pathway. The Plant Cell 13(6): 1293–1304.

Lu SX, Webb CJ, Knowles SM, Kim SHJ, Wang Z, Tobin EM. 2012. CCA1 and ELF3 Interact in the Control of Hypocotyl Length and Flowering Time in Arabidopsis. Plant Physiol 158(2): 1079–1088.

McWatters HG, Bastow RM, Hall A, Millar AJ. 2000. The ELF3zeitnehmer regulates light signalling to the circadian clock. Nature 408(6813): 716–720.

Millar AJ, Carre IA, Strayer CA, Chua N-H, Kay SA. 1995. Circadian clock mutants in Arabidopsis identified by luciferase imaging. Science 267(5201): 1161–1163.

Mizuno T, Nomoto Y, Oka H, Kitayama M, Takeuchi A, Tsubouchi M, Yamashino T. 2014. Ambient Temperature Signal Feeds into the Circadian Clock Transcriptional Circuitry Through the EC Night-Time Repressor in Arabidopsis thaliana. Plant and Cell Physiology 55(5): 958–976.

Nakamichi N, Kiba T, Henriques R, Mizuno T, Chua N-H, Sakakibara H. 2010. PSEUDO-RESPONSE REGULATORS 9, 7, and 5 Are Transcriptional Repressors in the Arabidopsis Circadian Clock. The Plant Cell Online 22(3): 594–605.

Nieto C, López-Salmerón V, Davière J-M, Prat S. 2015. ELF3-PIF4 interaction regulates plant growth independently of the Evening Complex. Current Biology 25(2): 187–193.

Niwa Y, Yamashino T, Mizuno T. 2009. The circadian clock regulates the photoperiodic response of hypocotyl elongation through a coincidence mechanism in Arabidopsis thaliana. Plant and Cell Physiology 50(4): 838–854.

Nohales MA, Kay SA. 2016. Molecular mechanisms at the core of the plant circadian oscillator. Nat Struct Mol Biol 23(12): 1061–1069.

Nozue K, Covington MF, Duek PD, Lorrain S, Fankhauser C, Harmer SL, Maloof JN. 2007. Rhythmic growth explained by coincidence between internal and external cues. Nature 448(7151): 358–361.

Nusinow DA, Helfer A, Hamilton EE, King JJ, Imaizumi T, Schultz TF, Farre EM, Kay SA. 2011. The ELF4-ELF3-LUX complex links the circadian clock to diurnal control of hypocotyl growth. Nature 475(7356): 398–402.

Oakenfull RJ, Davis SJ. 2017. Shining a light on the Arabidopsis circadian clock. Plant, Cell & Environment 40(11): 2571–2585.

Panigrahi KCS, Mishra P. 2015. GIGANTEA - An Emerging Story. Frontiers in Plant Science 6.

Park Y-J, Kim JY, Lee J-H, Lee B-D, Paek N-C, Park C-M. 2020. GIGANTEA Shapes the Photoperiodic Rhythms of Thermomorphogenic Growth in Arabidopsis. Molecular Plant 13(3): 459–470.

Quint M, Delker C, Franklin KA, Wigge PA, Halliday KJ, Zanten M. 2016. Molecular and genetic control of plant thermomorphogenesis. Nat Plants 2.

Raschke A, Ibanez C, Ullrich K, Anwer M, Becker S, Glockner A, Trenner J, Denk K, Saal B, Sun X, et al. 2015. Natural variants of ELF3 affect thermomorphogenesis by transcriptionally modulating PIF4-dependent auxin response genes. BMC Plant Biology 15(1): 197.

Reed JW, Nagpal P, Bastow RM, Solomon KS, Dowson-Day MJ, Elumalai RP, Millar AJ. 2000. Independent action of ELF3 and phyB to control hypocotyl elongation and flowering time. Plant Physiology 122(4): 1149–1160.

Ronald J, Davis S. 2017. Making the clock tick: the transcriptional landscape of the plant circadian clock.

Salomé PA, McClung CR. 2005. PSEUDO-RESPONSE REGULATOR 7 and 9 Are Partially Redundant Genes Essential for the Temperature Responsiveness of the Arabidopsis Circadian Clock. The Plant Cell Online 17(3): 791–803.

Salomé PA, Weigel D, McClung CR. 2010. The role of the Arabidopsis morning loop components CCA1, LHY, PRR7, and PRR9 in temperature compensation. The Plant Cell 22(11): 3650–3661.

Sanchez SE, Rugnone ML, Kay SA. 2020. Light Perception: A Matter of Time. Molecular Plant 13(3): 363–385.

Schaarschmidt S, Lawas LMF, Glaubitz U, Li X, Erban A, Kopka J, Jagadish SVK, Hincha DK, Zuther E. 2020. Season Affects Yield and Metabolic Profiles of Rice (Oryza sativa) under High Night Temperature Stress in the Field. Int J Mol Sci 21(9).

Thines B, Harmon FG. 2010. Ambient temperature response establishes ELF3 as a required component of the core Arabidopsis circadian clock. Proc Natl Acad Sci U S A 107(7): 3257–3262.

Tseng T-S, Salomé PA, McClung CR, Olszewski NE. 2004. SPINDLY and GIGANTEA interact and act in Arabidopsis thaliana pathways involved in light responses, flowering, and rhythms in cotyledon movements. The Plant Cell 16(6): 1550–1563.

Yamashino T, Ito S, Niwa Y, Kunihiro A, Nakamichi N, Mizuno T. 2008. Involvement of Arabidopsis Clock-Associated Pseudo-Response Regulators in Diurnal Oscillations of Gene Expression in the Presence of Environmental Time Cues. Plant and Cell Physiology 49(12): 1839–1850.

Zagotta MT, Hicks KA, Jacobs CI, Young JC, Hangarter RP, Meeks-Wagner DR. 1996. The Arabidopsis ELF3 gene regulates vegetative photomorphogenesis and the photoperiodic induction of flowering. The Plant Journal 10(4): 691–702.

Zielinski T, Moore AM, Troup E, Halliday KJ, Millar AJ. 2014. Strengths and limitations of period estimation methods for circadian data. PLOS ONE 9(5): e96462.

